# TIME-Seq Enables Scalable and Inexpensive Epigenetic Age Predictions

**DOI:** 10.1101/2021.10.25.465725

**Authors:** Patrick T Griffin, Alice E Kane, Alexandre Trapp, Jien Li, Matthew Arnold, Jesse R Poganik, Maeve S McNamara, Margarita V Meer, Noah Hoffman, João Amorim, Xiao Tian, Michael R MacArthur, Sarah J Mitchell, Amber L Mueller, Colleen Carmody, Daniel L Vera, Csaba Kerepesi, Nicole Noren Hooten, James R Mitchell, Michele K Evans, Vadim N Gladyshev, David A Sinclair

**Author notes:** Institute for Systems Biology, Seattle, WA 98109 USA.

## Abstract

Epigenetic “clocks” based on DNA methylation (DNAme) have emerged as the most robust and widely employed aging biomarkers, but conventional methods for applying them are expensive and laborious. Here, we develop <u>T</u>agmentation-based Indexing for <u>M</u>ethylation <u>Seq</u>uencing (TIME-Seq), a highly multiplexed and scalable method for low-cost epigenetic clocks. Using TIME-Seq, we applied multi-tissue and tissue-specific epigenetic clocks to over 1,600 mouse DNA samples. We also discovered a novel approach for age prediction from shallow sequencing (e.g., 10,000 reads) by adapting *scAge* for bulk measurements. In benchmarking experiments, TIME-Seq performed favorably against prevailing methods and could quantify the effects of interventions thought to accelerate, slow, and reverse aging in mice. Finally, we built and validated a highly accurate human blood clock from 1,056 demographically representative individuals. Our methods increase the scalability and reduce the cost of epigenetic age predictions by more than 100-fold, enabling accurate aging biomarkers to be applied in more large-scale animal and human studies.

## Introduction

Aging is difficult to study, in part, because it is difficult to quantify^1^. In recent years, researchers have attempted to address this problem with aging “clocks”, which are machine learning-derived biomarkers trained to predict age or age proxies^2^. Since clock predictions are not perfect, individuals are often predicted younger or older than their chronological age, and this difference is hypothesized to reflect variation in the biological rate of aging^3^. Both physiological measurements^4,5^ and biomolecules^3,6,7^ have been used to build aging clocks and they are becoming increasingly common readouts to assess longevity interventions. Despite their promise, accurate clocks that are inexpensive and easily applied to large studies are lacking.

The most robust and widely used aging clocks are based on DNA cytosine methylation (DNAme) and are interchangeably referred to as DNAme clocks or epigenetic clocks. These clocks comprise sets of CpGs and corresponding algorithms that use methylation levels to predict age. Epigenetic clocks have been built for humans^8-10^, mice^11-13^, and a multitude of other mammals^14-17^, and they have been shown to reflect interventions that are associated with longevity^18^, accelerated aging^19^, and even cellular rejuvenation^20,21^. While much focus has been put on developing more accurate clocks or clocks adjusted by health outcomes^10,22^, very little work has been done to make epigenetic clocks more experimentally tractable.

Ideally, epigenetic clocks could be measured for low-cost with a scalable technology. However, clocks are predominantly built and assayed using Illumina BeadChip^23^ microarrays or <u>R</u>educed <u>R</u>epresentation <u>B</u>isulfite <u>S</u>equencing^24^ (RRBS), which are laborious and cost hundreds of dollars per-sample. While these methods are useful for biomarker discovery because they measure hundreds-of-thousands to millions of CpGs, they are excessive for the accurate measurement of epigenetic clocks that typically only require several hundred loci to be measured. Thus, a more economical and targeted approach could be used for epigenetic clocks with similar accuracy and robustness.

With this in mind, we developed <u>T</u>agmentation-based Indexing for <u>Me</u>thylation <u>Seq</u>uencing (TIME-Seq), an optimized bisulfite-sequencing approach to enable low-cost and scalable epigenetic age predictions. We use TIME-Seq to build four new epigenetic clocks for mice and one clock for humans, applying them to 2772 unique samples from 9 different tissue and cell types. We benchmark TIME-Seq against the prevailing methods and validate our clocks in independent cohorts as well as interventions that alter the rate of aging. Using *scAge*, an algorithm originally applied to single-cell epigenetic age analysis^25^, we discover it is possible to accurately predict age from as few as 10,000 TIME-Seq reads^25^. Our methods decrease costs of epigenetic clock analysis by more than two orders of magnitude, promising to expedite their use in more large-scale experiments.

## Results

We designed TIME-Seq, a novel targeted sequencing method to build and measure DNAme-based biomarkers (e.g., epigenetic clocks) for low-cost in hundreds to thousands of samples. TIME-Seq leverages barcoded and sodium bisulfite-resistant Tn5-transposomes to rapidly index sample DNA for a pooled library preparation (Fig. 1a), which streamlines large-scale experiments and minimizes the cost of consumables (Supplementary Table 1). After tagmentation and pooling, methylated end-repair (5-methyl-dCTP replaces dCTP) is performed, and pools are prepared for in-solution hybridization enrichment using biotinylated-RNA baits (Extended Data Fig. 1a-e). Unlike bisulfite-compatible single-cell indexing approaches^26^, we designed barcoded TIME-Seq adaptors to be short (38-nt) for optimal enrichment efficiency since longer adaptors are more likely to daisy-chain with off-target DNA^27^ (Extended Data Fig. 1f-h). Baits are produced in-house from single-stranded oligonucleotide libraries (Supplementary Table 2), providing inexpensive enrichments from a regenerable source. After bisulfite conversion of captured DNA and indexed-PCR amplification of each pool, Illumina short-read sequencing is performed (Extended Data Fig. 2) and sample reads are demultiplexed based on pool and Tn5-adaptor indexes. From mapped reads, a matrix of methylation values for CpGs in each sample is used to train or predict a DNAme biomarker.

**Figure 1.**
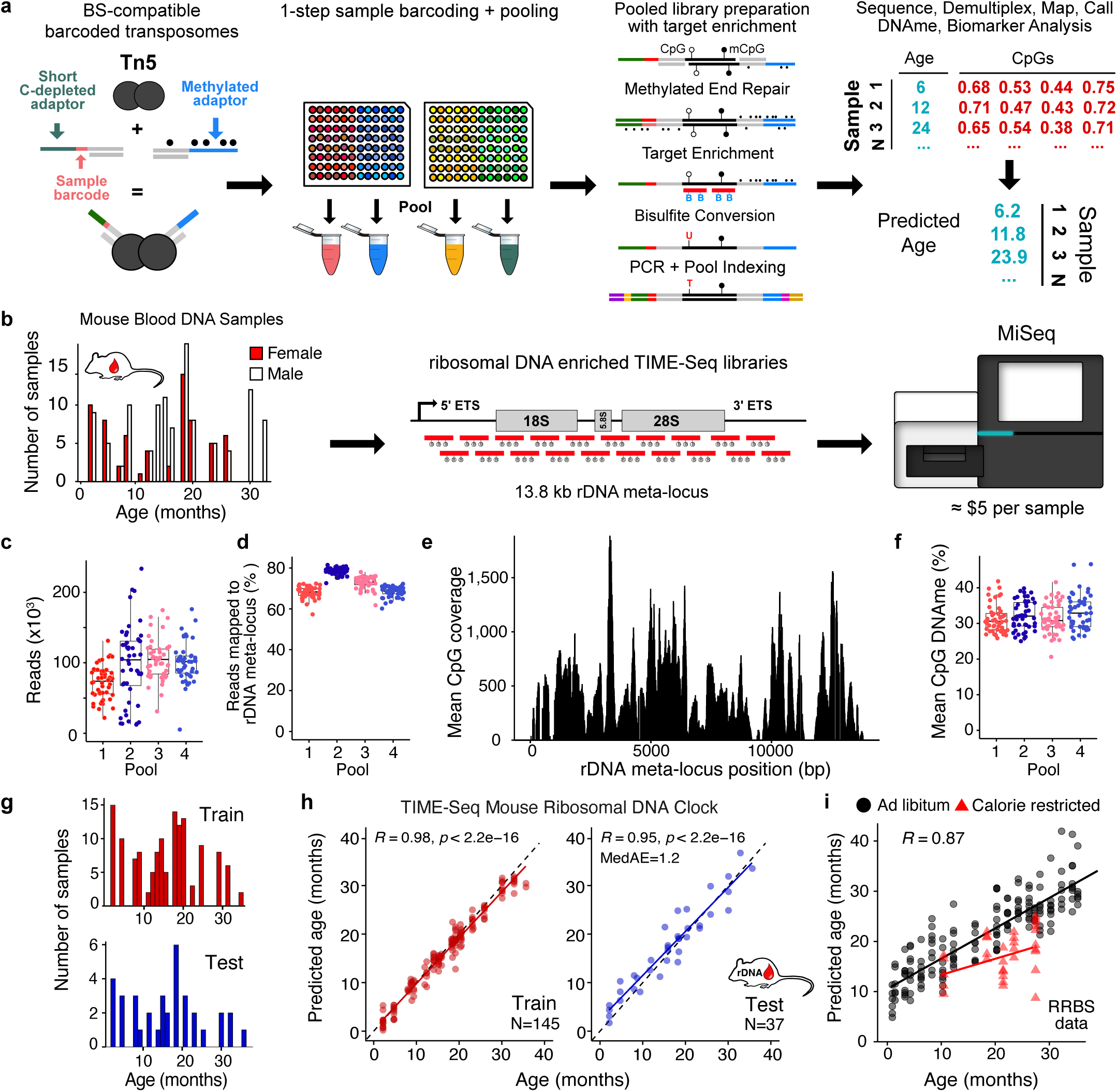
TIME-Seq enables highly efficient epigenetic age predictions. **a**, Schematic of the TIME-Seq library preparation for highly multiplexed targeted methylation sequencing to build and measure DNA methylation (DNAme)-based biomarkers. **b**, Proof-of-concept rDNA clock experiment schematic. 191 mouse blood DNA samples (histogram) were prepared with TIME-Seq enriched for rDNA and sequenced for clock training and testing. **c**, Reads demultiplexed from each rDNA clock pool of 47-48 samples. **d**, Percent of demultiplexed reads from each sample that mapped to the rDNA meta-locus. **e**, Mean coverage at rDNA meta-locus CpGs in rDNA-enriched TIME-Seq libraries. **f**, Mean CpG methylation from each sample in the four pools. **g**, Histogram of training (N=145; red) and testing (N=37; blue) samples used to develop the TIME-Seq rDNA clock. **h**, TIME-Seq mouse rDNA clock showing age prediction for training (red) and testing (blue). Pearson correlation and median absolute error (MedAE) on test are shown in the top left corner. **i**, TIME-Seq-based clock developed using only CpGs with at least 50 coverage in RRBS data used to develop the original mouse rDNA clock. Caloric restricted (CR) mice are represented as red triangles.

Ribosomal DNA (rDNA) is a highly repetitive locus that shows increased DNAme with age in mice, allowing for accurate epigenetic clocks to be constructed^21,28^. Therefore, we performed a small-scale pilot of TIME-Seq in C57BL/6 mouse blood DNA samples using hybridization probes against the described rDNA clock CpGs^28^ (Extended Data Fig. 3a). Samples efficiently demultiplexed from each TIME-Seq pool, and DNA methylation was accurately measured (Extended Data Fig. 3b-c) with a high correlation (*R*≥0.90) between replicate CpG levels and deep coverage at targeted epigenetic clock loci from less than 600,000 reads (Extended Data Fig. 3d-f). Compared to RRBS libraries of the same samples, TIME-Seq libraries had substantially higher overlap with target clock CpGs (Extended Data Fig. 3g). Age prediction using an existing RRBS-based rDNA clock, however, showed only moderate correlation with age in our pilot (R=0.53; Extended Data Fig. 3h), possibly due to the differences in CpG coverage between TIME-Seq and RRBS at several clock loci (Extended Data Fig. 3i).

To build a more accurate rDNA clock compatible with TIME-Seq, enrichment baits tiling the entire rRNA promoter and coding regions were designed and used to enrich TIME-Seq libraries from 191 C57BL/6 mouse blood DNA samples (ages 2-35 months) in pools of 47-48 (Fig. 1b). Pools were combined and sequenced on an Illumina MiSeq for a per-sample cost of less than five US dollars (USD) (see Supplementary Table 3 for sequencing costs). The majority of demultiplexed reads (Fig. 1c) from each sample mapped to the rDNA repeat meta-locus (Fig. 1d), resulting in high coverage for each sample at rDNA CpGs (Fig. 1e) and accurate DNA methylation quantification (Fig. 1f).

To train an epigenetic clock, the 182 samples that passed quality filters were split approximately 80:20 into training, and testing sets (Fig. 1g) and elastic net regression (α=0.05) was applied to the training data. After fine-tuning our model on the training set (see Methods), age predictions using the resulting 232 CpG TIME-Seq rDNA Clock showed a high correlation with age (training, *R*=0.98; testing, *R*=0.95) and a median absolute error (MedAE) of only 1.2 months in the testing samples (Fig. 1h). To build a clock that could be applied to both TIME-Seq and RRBS, we trained a model from TIME-Seq data using only CpGs with high coverage in an existing RRBS data set^11^. This clock showed a high age correlation (*R*=0.87) when applied to RRBS data and reflected the longevity benefit of caloric restriction (Fig. 1i).

In contrast to rDNA clocks, those built with CpGs from across the genome reflect diverse changes in aging hallmarks and their associated genes^3^. Therefore, we designed hybridization enrichment probes for 957 distinct CpG islands in gene promoters or other gene regulatory elements previously reported to have high age-correlation in mouse blood^11^ and multi-tissue clocks^12,13^ (Extended Data Fig. 4a). With these enrichment probes, we prepared TIME-Seq libraries using 1019 DNA samples from mouse blood, liver, skin, kidney, white adipose tissue (WAT), or muscle, and sequenced libraries for an average per-sample cost of $5.21. From the data, we prepared a high-coverage methylation matrix of 6373 CpGs for clock training (Fig. 2a). Variation between samples was mainly due to the tissue of origin (Fig. 2b) as expected from quality methylation sequencing datasets from multiple tissues.

**Figure 2.**
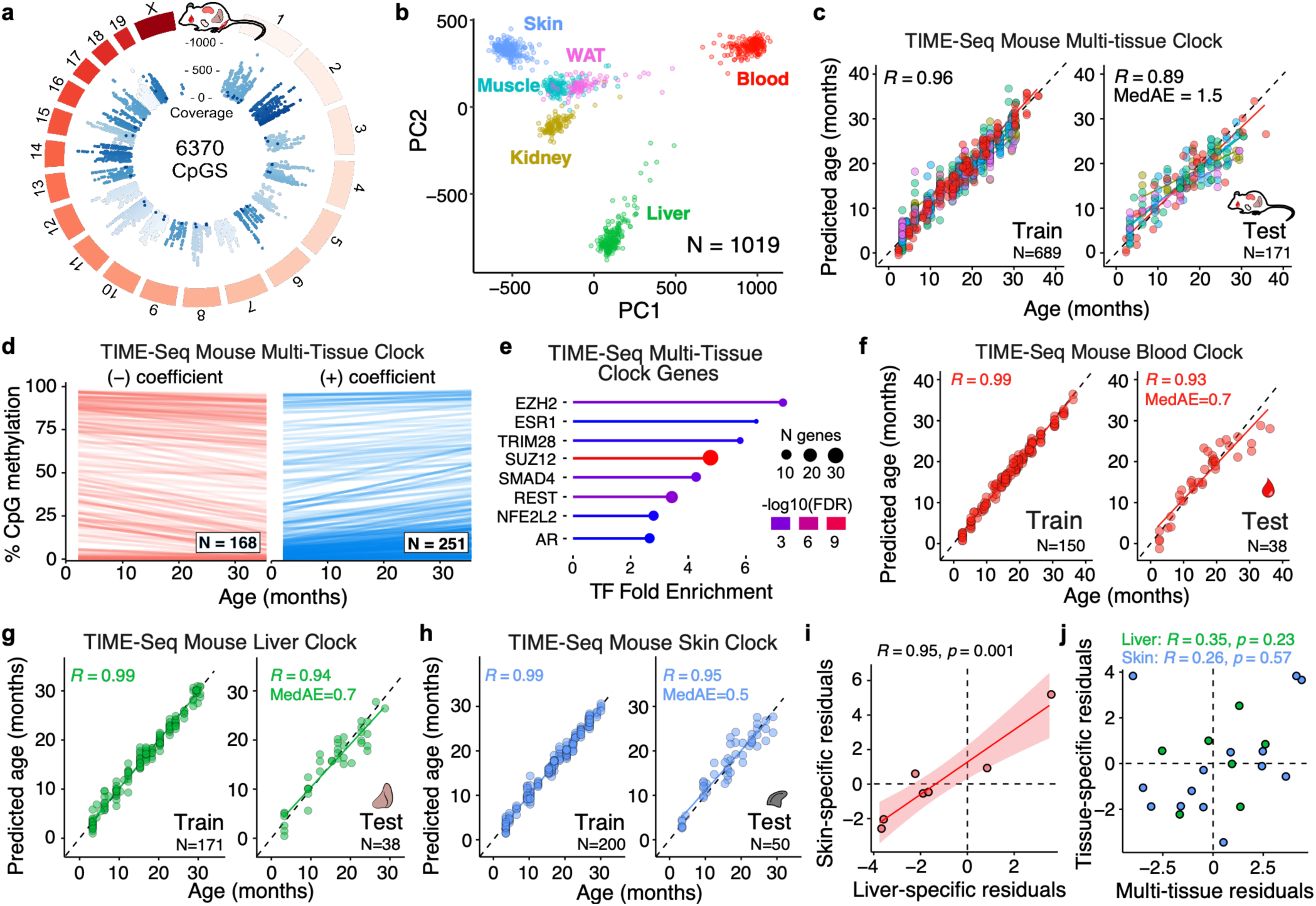
Low-cost TIME-Seq multi-tissue and tissue-specific clocks applied to 1019 mouse tissue samples. **a**, Circular genome plot illustrating the position and mean coverage of the 6370 high coverage CpGs from TIME-Seq libraries in 1019 mouse tissue samples. **b**, Principal component analysis for TIME-Seq data colored by their tissue-of-origin. Liver (green), blood (red), skin (blue), muscle (turquoise), Kidney (yellow), and white adipose tissue (abbreviated WAT; hot pink). **c**, TIME-Seq Mouse Multi-tissue Clock train (left) and test (right) predictions plotted against chronological age. **d**, DNA methylation change with age in TIME-Seq Mouse Multi-tissue CpGs split by their clock coefficient sign. The absolute value of the clock coefficient is coded in the transparency of each line. **e**, Enrichment of transcription factor binding sites from genes associated with the TIME-Seq Mouse Multi-tissue Clock. **f-h**, TIME-Seq Mouse Blood Clock (**f**), Liver Clock (**g**); and Skin Clock (**h**) with train (left) and test (right) predictions plotted against chronological age. **i**, Age-adjusted residuals for Liver and Skin Clocks from the same mice predicted in the testing sets. **j**, Age-adjusted residuals for either Skin or Liver Clocks plotted against the predictions from the Multi-tissue Clock in the same sample.

Next, we trained and tested the TIME-Seq Mouse Multi-tissue clock, which accurately predicts age in mouse blood, liver, skin, kidney, and WAT (Testing *R*= 0.89; MedAE = 1.5 months). As described in previous studies^8,21^, when using age-correlated CpGs from other tissues, the prediction of age in muscle was less accurate (Extended Data Fig. 4b-c) and this tissue was excluded from multi-tissue clock training. Our 419-CpG Mouse Multi-Tissue Clock contains both positive and negative age-correlated CpGs (Fig. 2d) that are enriched for genes regulated by PRC2 components (e.g., EZH2 and SUZ12) and the longevity-associated transcription factor REST^29^ (Fig. 2e). We also trained tissue-specific epigenetic clocks (Fig. 2f-h) for mouse blood (testing: *R* = 0.93, MedAE = 0.7 months), liver (testing: *R*=0.94, MedAE = 0.7 months), and skin (testing: *R* = 0.95, MedAE = 0.5 months). The TIME-Seq Mouse Blood Clock is enriched for genes associated with regulating pluripotency of stem cells, while all three tissue-specific clocks contained enrichment of genes regulated by PRC2 components, as well as tissue-specific TFs associated with clock genes (Extended Data Fig. 4d-g).

It is still largely unknown what factors influence epigenetic clocks and how different aging clocks relate to each other^2^. It has been hypothesized that certain clocks exhibit more environmental “extrinsic” influence, whereas other clock are more intrinsically defined (i.e., influenced more by genetic variation)^3^. To understand how our clocks relate to each other, we identified mice that contributed tissues to 2 or more testing sets in clock development. To control for slight bias in prediction error with age, we calculated the age-adjusted prediction residuals for each clock (see Methods) and plotted the pairwise data for each tissue and clock (Fig. 2i-j). There was highly positive correlation (*R*=0.95, p=0.001) for the TIME-Seq Skin and TIME-Seq Liver Clocks with each other, but no significant correlation for either tissue-specific clock with the TIME-Seq Multi-tissue Clock. This result suggests there may be some extrinsic factor contributing to synchrony between the TIME-Seq Liver and Skin Clocks, whereas our Multi-tissue Clock is not subject to the same influence and reflects other aging modules.

Epigenetic clocks rely on ratios of methylated to unmethylated CpGs generated from dozens or hundreds of DNA molecules at each clock locus. For sequencing-based clocks, this necessitates expensive deep sequencing. Recently, the probabilistic age-prediction algorithm *scAg*e was introduced to overcome this constraint and make accurate predictions from semi-binary single-cell DNAme data^25^. We hypothesized that *scAge* could be used for accurate age prediction from shallow sequencing in bulk DNA samples (e.g., 1/100^th^ depth), which would resemble sparse DNAme data from single cells. To test this, we performed low-pass sequencing on three TIME-Seq pools containing 121 mouse-blood DNA samples (Fig. 3a; 119 passed quality filters). Sample reads ranged from 3,610 to 32,588 (median 11,560) with a per-sample cost of just $1.85 (Fig. 3b). While average methylation levels were approximately the same as deep-sequenced data (Extended Data Fig. 5a), the vast majority of CpGs were only covered by one or a few reads (Fig. 3c).

**Figure 3.**
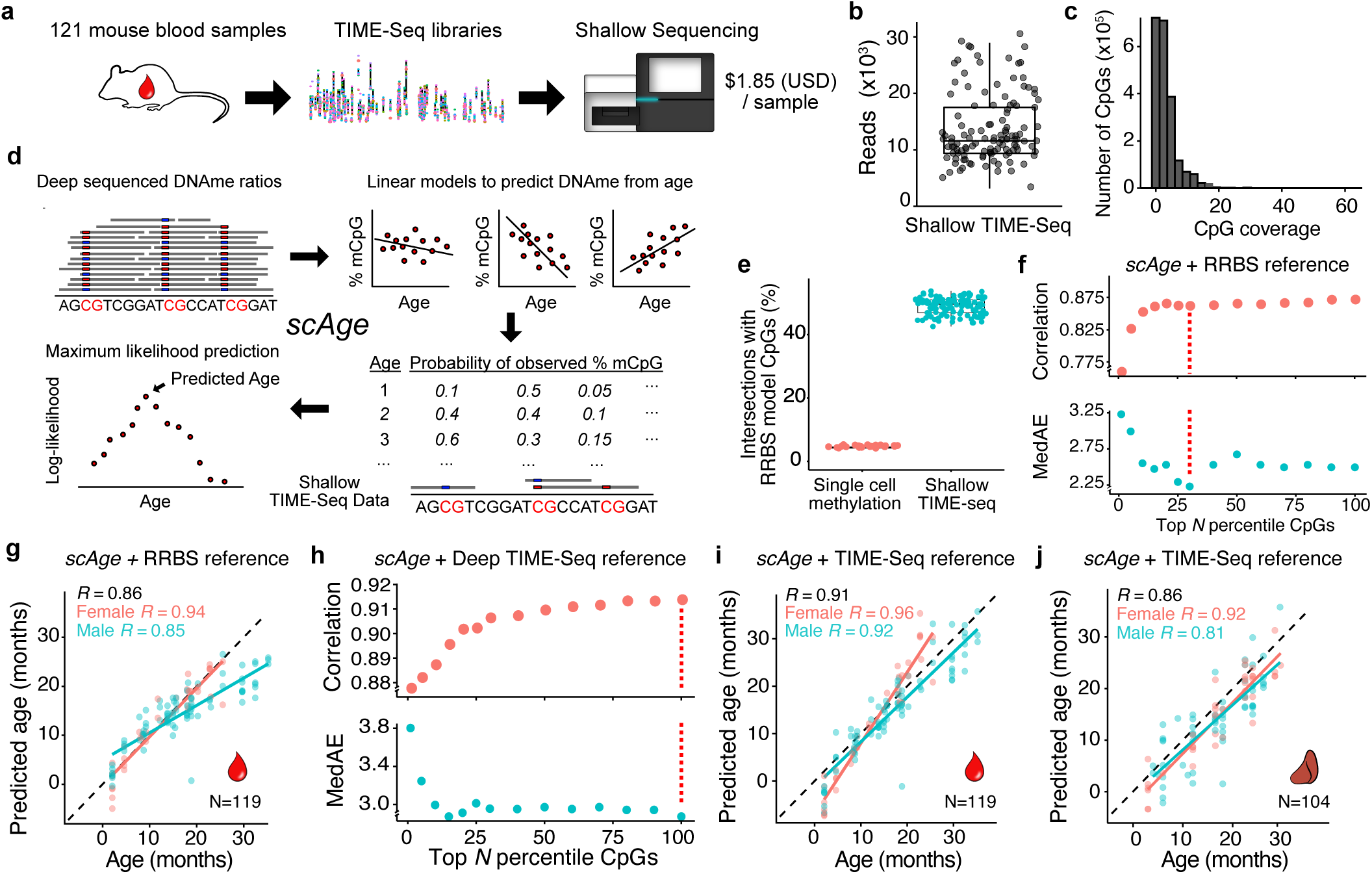
Ultra-cheap age prediction with scAge from shallow sequencing of TIME-Seq libraries. **a**, Schematic of shallow sequencing of 121 MD-enriched TIME-Seq libraries. **b**, Demultiplexed read number from each shallow sequenced sample. **c**, Histogram of CpG coverages in shallow sequenced samples. **d**, Schematic of the scAge framework for maximum likelihood age prediction from shallow TIME-Seq methylation data. **e**, Percent of scAge model CpGs covered in each shallow sequenced TIME-Seq library and a previously reported single-cell methylation dataset, Gravina et al., (2016). **f**, Pearson correlation and MedAE from scAge-based age prediction using Thompson et al., (2018) reference RRBS data. CpGs in each sample were ranked by the absolute value of their correlation with age (based on deeply sequenced data), and only the top N percentile was used for maximum likelihood prediction at each point. The red line indicates the percentile presented in (g). **g**, Age predictions from scAge in shallow-sequenced TIME-Seq libraries (N=119) using RRBS reference data (top 30% age-associated CpGs). Pearson correlations are shown in the top left corner. **h**, Pearson correlation and MedAE from scAge-based age prediction of the top N percentiles of CpGs using deep-sequenced, MD-enriched TIME-Seq data as reference (data used for Fig. 2k-o). The red line indicates the percentile presented in (i). **i**, scAge predictions from shallow-TIME-Seq data using deep-sequenced TIME-Seq data (100% of CpGs) as models. Pearson correlations are shown in the top left corner. **j**, scAge predictions from 104 shallow-sequenced liver samples using deep-sequenced TIME-Seq liver data as model CpGs (100% of CpGs). Pearson correlations are shown in the top left corner.

The *scAge* algorithm leverages existing deep-sequenced methylation data to construct linear models for maximum-likelihood age prediction from sparse data (Fig. 3d). We modified *scAge* to allow for more than just semi-binary data (see Methods) and applied it to predict age in our shallow-TIME-Seq experiment using published RRBS data^11,13^ as reference. Since TIME-Seq libraries are enriched for age-correlated loci in RRBS datasets, there was high intersection (median 49.1%) between model CpGs and shallow-TIME-Seq data, compared to a more random distribution of sparse data, such as whole-genome single-cell methylation data^30^ (Fig. 3e).

In support of our hypothesis, age predictions from *scAge* in shallow-TIME-Seq data were highly accurate (Fig. 3f-g and Extended Data Fig. 5b-c) with a correlation of *R*=0.94 for females, *R*=0.85 for males, and a MedAE of just 2.23 months for all predictions. Next, we used the deep-sequenced TIME-Seq data as the reference for *scAge* (Fig. 3h-i), which further improved age correlation in the entire dataset to *R=*0.91 with a MedAE of 2.87 months. To test if this approach is generalizable to other mouse tissues, we prepared and shallow-sequenced TIME-Seq libraries from 104 mouse liver samples (ages 3-29 months). Using both deep-sequenced TIME-Seq liver data (Fig. 3j and Extended Data Fig. 5d) and RRBS liver data (Extended Data Fig. 5e-f) as a reference, age predictions were accurate.

To validate the robustness of TIME-Seq-based age predictions, we prepared TIME-Seq libraries in two separate experiments using blood DNA from an independent cohort of mice (Fig. 4a). A subset of these mice had been tracked longitudinally, assessed using the mouse frailty-index^31^, and had blood composition parameters measured. The TIME-Seq Blood Clock (*R*=0.93; Fig. 4b), the TIME-Seq Multi-tissue Clock (*R*=0.91; Fig. 4c), and the combination of *scAge* and shallow-TIME-Seq (*R*=0.87; Fig. 4d and Extended Data Fig. 6a) provided accurate predictions that reflected aging longitudinally. For TIME-Seq libraries enriched for rDNA (Extended Data Fig. 6b-c), mice from the same colony as the original training set validated clock rDNA clock accuracy (JAX mice, *R=*0.96). As expected, mice from different colonies gave more variable predictions (NIA mice, *R* = 0.81) since rDNA copy number varies greatly between different mouse strains, colonies, and even individuals within a colony^32^ and copy number directly relates to aggregated methylation status^33^.

**Figure 4.**
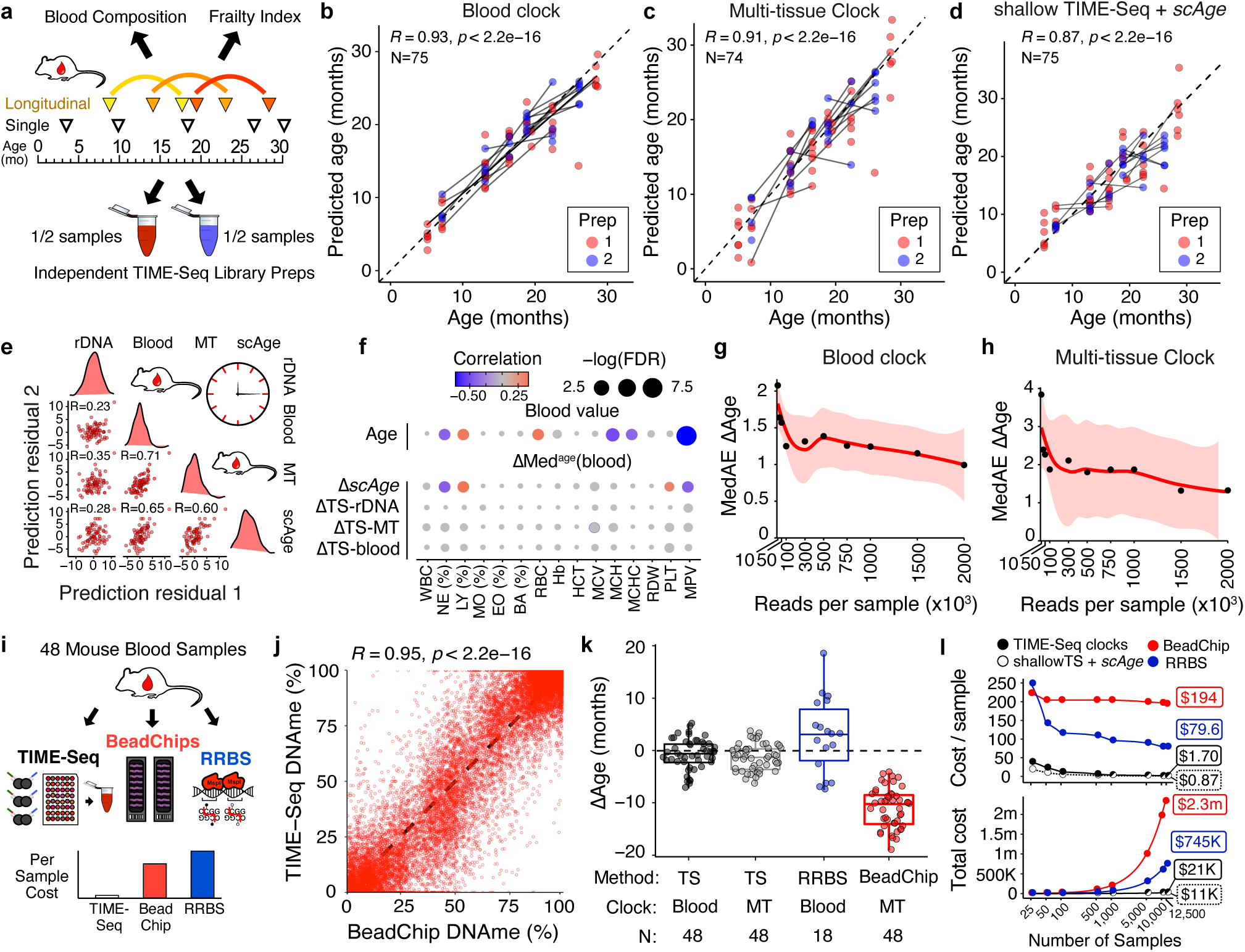
TIME-Seq is a robust and scalable alternative to conventional clock approaches. **a**, Experimental schematic for validation of TIME-Seq age prediction methods in an independent cohort of mice with longitudinal timepoints, paired frailty index, and blood composition data. **b-d**, TIME-Seq age predictions in two independent validation library preparations using (**b**) the TIME-Seq Mouse Blood Clock, (**c**) the TIME-Seq Mouse Multi-tissue Clock, and (**d**) scAge predictions from shallow-TIME-Seq data (shallow-TS + scAge) using deep-TIME-Seq data as model CpGs. Lines connect the same mouse at two different ages. Pearson correlations and n values are shown in the top left corner. **e**, Correlation between age-adjusted prediction residuals in the validation sets from the different prediction approaches. **f**, Correlation and significance matrix between ΔAge from each approach and ΔMed^age^(blood), i.e., the difference in median value from similar aged mice for each blood measurement. Color and size of each circle represent the correlation and p-value significance, respectively. WBC = white blood cell count, NE (%) = percent of neutrophils, LY (%) = percent of lymphocytes, MO (%) = percent of monocytes, EO (%) = percent of eosinophils, BA (%) = percent of basophils, RBC = red blood cell count, Hb = hemoglobin, HCT = hematocrit, MCV = mean corpuscular volume, MCH = mean corpuscular hemoglobin, MCHC = mean corpuscular hemoglobin concentration, RDW = red blood cell distribution width, PLT = platelets, MPV = mean platelet volume. **g-h**, Sequencing saturation simulation to estimate clock accuracy from different read numbers in the validation samples. **i**, Schematic of benchmarking experiment to compare TIME-Seq to Illumina BeadChip and RRBS. **j**, Comparison of CpG methylation percent in TIME-Seq and BeadChip. Each dot represents the same CpG from the same sample measured each technology. **k**, Comparison of ΔAges for each method and the associated clocks. **i**, Comparison of cost per sample (top) and total cost (bottom) for TIME-Seq clocks (black outline; black fill; black solid line), shallow TIME-Seq and scAge (black outline; white fill; dashed line), BeadChip (red), and RRBS (blue) across a range of sample scales. Half-filled circles denote points that are overlapping between TIME-Seq clocks and scAge. Million is abbreviate as “m”. “K” denotes thousand.

To understand if our age prediction methods were synchronous in predicting animals older or younger, we plotted the pairwise age-adjusted prediction residuals for each mouse from each method. While all prediction methods were significantly positively correlated, the TIME-Seq Multi-Tissue Clock, Blood Clock, and *scAge* age-adjusted prediction residuals were much more highly correlated with each other and lowly correlated rDNA clock residuals (Fig. 4e), suggesting rDNA methylation may be capturing a separate aging module only partially reflected in the other clock methods. The high correlation between Blood and Multi-tissue Clock prediction residuals suggests that similar aging modules are reflected by these clocks, in contrast with Multi-tissue Clock asynchrony with the Skin and Liver Clocks.

The difference between predicted age from epigenetic clocks and chronological age (ΔAge) has been shown to correlate with a wide variety of age-associated phenotypes^3^. To test if TIME-Seq predictions related to other measures of health or aging, we compared ΔAges from each approach to mouse frailty index and blood composition measurements, which have been shown to influence epigenetic clocks^2^. To control for the raw age correlation of each variable (top Fig. 4f and Extended Data Fig. 6d), measurements from each mouse were subtracted by the median value of that variable in similar aged animals—abbreviated ΔMed^age^(blood) and ΔMed^age^(FI). ΔMed^age^(blood) values were not correlated with ΔAges from the deep-sequenced clocks, suggesting blood cell composition was not driving predictive variance (Fig. 4f). In contrast, ΔMed^age^(platelet) values, as well as parameters with significant age correlations, mean platelet volume (MPV), percent neutrophils (NE %), and percent lymphocytes (LY %) were also significantly correlated with *scAge* ΔAges in the same direction, suggesting *scAge* predictions may be more susceptible to the influence of changes in aging blood composition. Since frailty index is also highly correlated with age and indicative of age-related decline, we assessed if ΔAges were correlated with ΔMed^age^(FI) values (i.e., whether mice that are frailer for their age are also predicted older and vice versa). Comparing ΔMed^age^(FI) to ΔAges (Extended Data Fig. 6e), we found no correlation with any TIME-Seq prediction method. This finding mirrors the previously described lack of correlation between frailty and human epigenetic age predictions from blood DNA (e.g., Hannum or Horvath clocks^34^) and suggests that frailty scores are a relatively distinct biomarker of health and functional decline.

To determine the minimum read number for accurate TIME-Seq clock prediction, we simulated a sequencing saturation experiment by extracting reads from each sample at a lower threshold, re-mapping the subset reads, and predicting age with the lower coverage (Fig. 4g-h). For the TIME-Seq Blood and Multi-tissue Clocks, prediction accuracy began to substantially decline with less than 500,000 reads per samples.

Next, we benchmarked TIME-Seq against the most common technologies for age prediction, Illumina Methylation BeadChip and RRBS, in 48 independent mouse blood samples (Fig. 4i). TIME-Seq methylation levels from the same CpG and same mouse were highly correlated with both BeadChip (*R*=0.95; Fig. 4j) and RRBS (*R*=0.92; Extended Data Fig. 7f). Epigenetic age predictions were highly accurate using either the TIME-Seq Blood or Multi-tissue Clocks, providing further independent validation (Fig. 4k). In contrast, the recently described BeadChip multi-tissue epigenetic clock^35^—the only mouse microarray-based clock that is commercially available—uniformly underpredicted the age of samples, possibly owing to a skewed age and sample distribution in the original study^35^. Finally, the RRBS-based mouse blood clock^11^ predicted samples with more accuracy than the BeadChip clock but with more variation than TIME-Seq.

To compare the cost of small- and large-scale clock analyses, we estimated costs for a range of sample sizes (Fig. 4i and Supplementary Table 5). TIME-Seq becomes increasingly cost-efficient at scale, with an estimated per-sample cost of just $1.70 for 12,500 samples. RRBS is estimated to be more expensive than Illumina BeadChip in small experiments but—also leveraging the efficiency of short-read sequencing—it becomes increasingly cheaper than BeadChip in large-scale application. At the largest scales, however, TIME-Seq is still approximately 50-fold less expensive than RRBS and 100-fold less than Illumina BeadChip-based analyses. Costs are even further minimized to just 87¢ per-sample if shallow sequencing of TIME-Seq followed by *scAge* is used. These results suggest that TIME-Seq clocks are a scalable and inexpensive alternative to more conventional methodologies.

Our goal for developing TIME-Seq was to apply clocks to large-scale intervention experiments such as those obtained in longitudinal mouse aging studies, *in vitro* screens, or large-scale human clinical trials. To understand if TIME-Seq clocks could detect differences in aging after interventions, we applied our clocks to controlled treatments associated with age deceleration, acceleration, or rejuvenation. We also applied TIME-Seq to an *in vitro* time-course to understand if it could be used for screening experiments.

Two of the most robust interventions that extend mouse healthspan and lifespan are amino acid restriction and caloric restriction^36^. Conversely, a reduction in healthspan and lifespan is seen when mice are fed a high fat diet (HFD)^37^. Using our TIME-Seq Mouse Blood Clock, mice that were 40% caloric restricted (CR) or methionine restricted (MetR) from age 24-months to 30-months were predicted younger than their *ad libitum* controls (Fig. 5a-b; MetR, p=0.007; 40% CR, p=0.021). Likewise, the TIME-Seq Liver Clock predicted that the livers of mice that experienced 30% caloric restriction for 10 months were younger (p = 0.046) than their *ad libitum* controls (Fig. 5c-d), with the TIME-Seq Multi-tissue Clock showing a trend towards younger prediction (p=0.053; Extended Data Fig. 7a). High-fat diet-fed mice were predicted older than mice of the same age on a standard diet using the TIME-Seq Liver Clock (p=0.026) and the TIME-Seq Multi-tissue Clock (p=0.018,), suggesting that age acceleration is reflected in our clock predictions. These data suggest that TIME-Seq clocks can detect age deceleration and acceleration from multiple tissues, even when interventions are initiated late in life.

**Figure 5.**
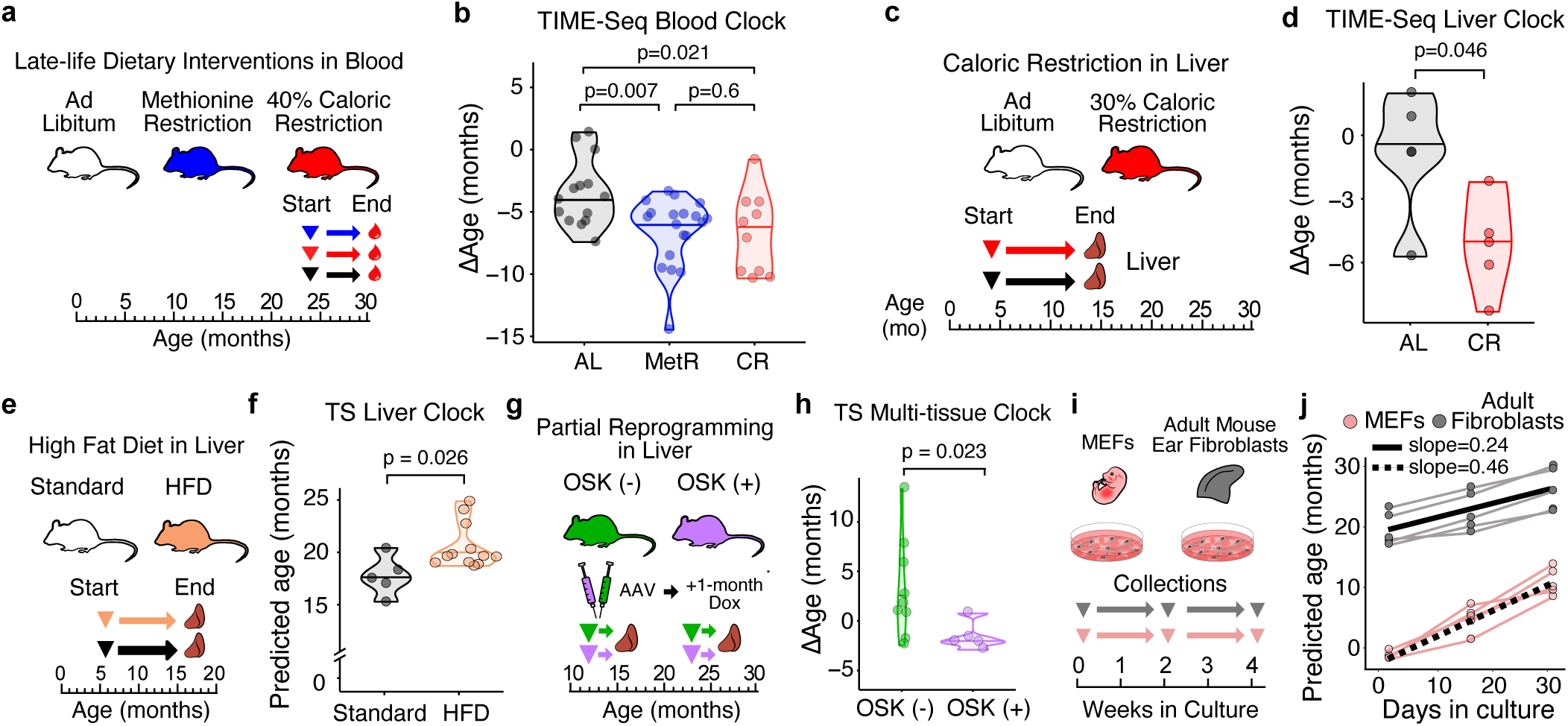
TIME-Seq clocks reflect interventions that slow, accelerate, and reverse aging and can be used for in vitro studies. **a**, Schematic of dietary restriction treatments started in late life. Blood was collected after 6-months of treatment. **b**, Comparison of ΔAges from TIME-Seq Blood Clock predictions in blood of dietary restricted and ad libitum mice. **c**, Schematic of caloric restriction started in young mice (4-months). Mice were treated for 10 months, and livers were collected. **d**, Comparison of ΔAge from ad libitum and CR mice using the TIME-Seq Mouse Liver Clock. **e**, Schematic of HFD treatment and liver collection. Mice were treated for 10 months starting at 6 months. **f**, Comparison of predicted ages for standard and high fat diet mice using the TIME-Seq Liver Clock. **g**, Schematic of AAV treatments with OSK-expressing or control (GFP) cassettes. Livers were collected 1-month after AAV injection. One set of control (N=4) and treatment (N=2) mice were 12 months, while another set was 24-months (control, N=5; treatment, N=3). **h**, Comparison of ΔAges in the livers of mice with (OSK+) or without (OSK-) OSK expression for 1-month. **i**, Schematic of experiment to assess MEFs and mouse adult ear fibroblasts in cell culture time course lasting 1 month with collection every 2-weeks. **j**, TIME-Seq multi-tissue clock predictions from cell culture samples collected across the time course.

Epigenetic reprogramming can rejuvenate aged tissues, driving gene expression and epigenetic changes toward a more youthful and regenerative state^20,38,39^. To understand if our clocks detected rejuvenation, we assayed mouse livers treated for 1-month with an AAV expressing an Oct-4, Sox-2, and Klf-4 (OSK) polycistronic gene cassette (Fig. 5g-h). With the TIME-Seq Multi-tissue Clock, mice with OSK expression were predicted significantly younger than control AAV-injected mice (p = 0.023). TIME-Seq Liver Clock predictions were not significantly different between groups (Extended Data Fig. 7c), perhaps reflecting differences between intrinsic and extrinsic aging modules. These results suggest that our method can detect epigenetic rejuvenation *in vivo*.

Epigenetic clocks have been shown to “tick” as cells grow in culture^19,40,41^, providing a promising way to screen for aging interventions *in vitro*. To test if TIME-Seq clocks also work on cultured cells, we grew five independent lines of low-passage mouse embryonic fibroblasts (MEFs) or adult mouse ear fibroblasts for one month and collected cells at three different timepoints (Fig. 5i). Using the TIME-Seq Multi-tissue Clock, MEFs were initial predicted a sub-zero (embryonic) age and steadily aged at a rate of approximately two weeks for every day in culture (Fig. 5j). Adult fibroblast lines were initial predicted 17-23 months and then aged at approximately half the rate as MEFs. Results were similar using the TIME-Seq Skin Clock, albeit with cells predicted to age at a slower rate (Extended Data 7d). These data indicate we can use TIME-Seq to track aging in cultured cells and provide further evidence for a different rate of aging between adult and embryonic cells^19^.

Most human clock studies train their age-prediction models on publicly available microarray data from dozens of past studies^8,10,19^ since new, large-scale experiments are exorbitantly expensive and laborious using microarrays. To develop a TIME-seq epigenetic clock that could be used in new large-scale human studies, we obtained 1056 human blood DNA samples from demographically representative individuals aged 18 to 103 years old (Fig. 6a). Most of the samples were designated for initial clock training and testing (N=796), whereas a subset was used for independent sample preparation and validation (N=260). Using enrichment probes against age-correlated loci from 11 described clocks^42^, TIME-Seq libraries were prepared and sequenced for a per-sample cost of $6.24. We achieved high coverage from clock CpGs across the genome (Fig. 6b), and sample data largely separated by age in the first two principal components (Fig. 6c), validating that age associated CpGs were enriched in the dataset.

**Figure 6.**
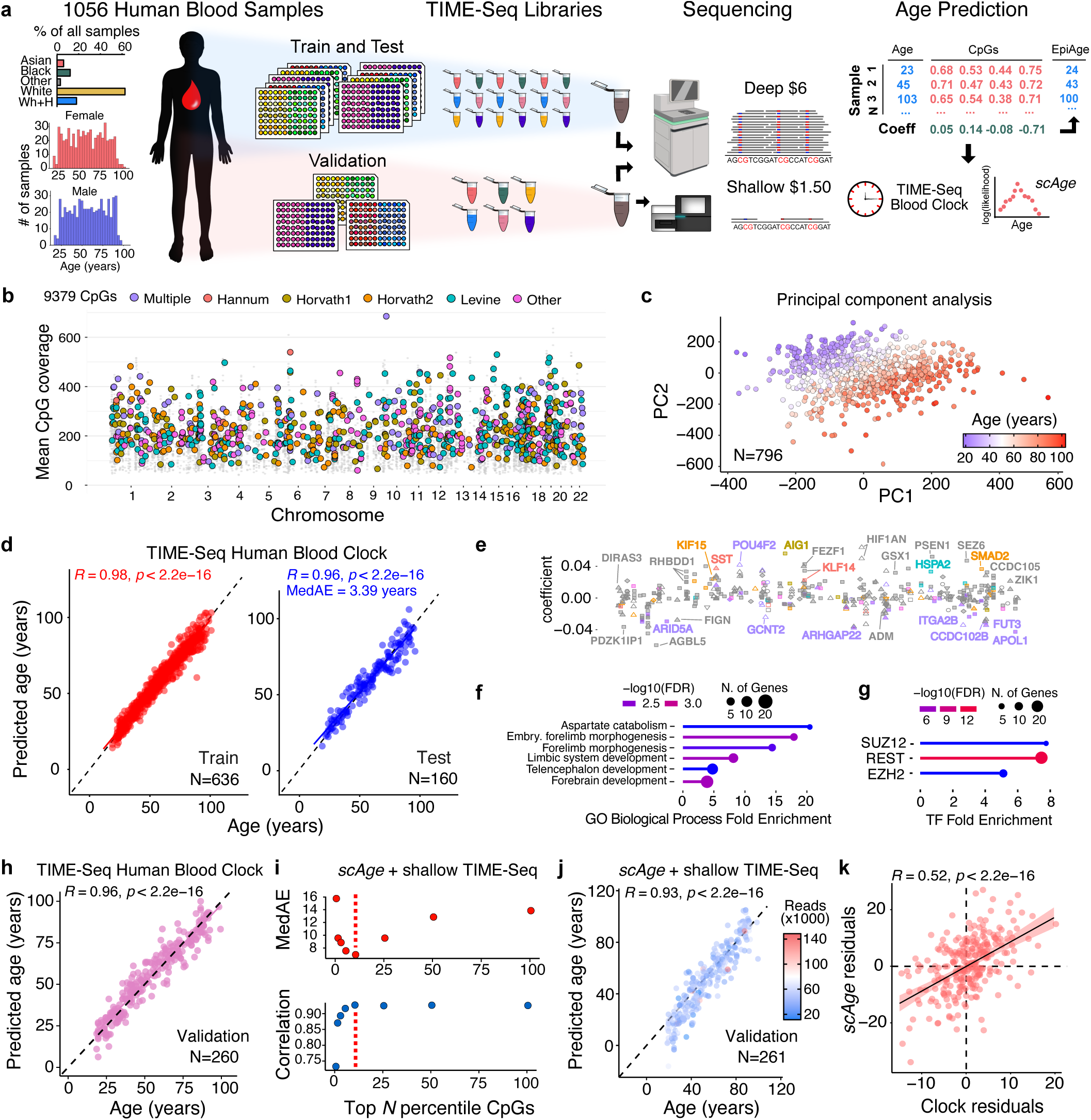
Highly accurate epigenetic age predictions in 1056 human blood samples using TIME-Seq clocks and scAge. **a**, Schematic of the experimental design to train, test, and validate TIME-Seq in 1056 human blood DNA samples. **b**, Coverage of 9379 CpGs from across the human genome. Colored dots are CpGs from described Illumina BeadChip clocks, whereas smaller grey dots are the other enriched CpGs. **c**, Principal component analysis of the methylation matrix for the train and test samples, colored by age from youngest (blue) to oldest (red). **d**, Predicted ages from the TIME-Seq Human Blood Epigenetic Clock in the train (right and test (left) samples. **e**, Annotation of the 405 clock CpGs with coefficient on the y-axis. X-axis (not shown) is genomic space in the same style as panel b from left (chromosome 1) to right (chromosome 22). CpGs and gene names are colored with the same color key as panel b. Feature annotation is coded by shape as follows: ○ 5’ UTR, ♦ exon, ▵ intergenic, ▴ intron, ▾ non-coding, ▪ promoter, and ▿ transcription termination site. **f-g**, Gene ontology (GO) analysis for enrichment of biological processes (**f**) or transcription factor (TF) binding sites (**g**) in genes associated with clock CpGs. **h**, TIME-Seq Human Blood Clock predictions in 260 independently prepared human blood DNA samples. **i**, Metrics for age prediction using various CpG percentiles from age-correlation ranking. Red line indicates the most accurate predictions shown in the next panel. **j**, Age predictions with scAge from shallow sequencing of TIME-Seq libraries using the top 10% of age-correlated CpGs from the original dataset. **k**, Age-adjusted residuals from clock and scAge predictions for validation samples.

With this data, we trained and tested the TIME-Seq Human Blood Clock (Fig. 6d), observing high age correlation in training (*R*=0.98) and testing predictions (*R*=0.96) with a MedAE of just 3.39 years, comparable to the most widely used human epigenetic clocks^8,9^. Gene set enrichment analysis of the resulting 405 CpG clock (Fig. 6e) revealed an association with developmental biological processes (Fig. 6f), including genes enriched for PRC2-associated proteins and REST similar to our mouse clocks (Fig. 6g).

To validate TIME-Seq in an independent experiment, we prepared and sequenced TIME-Seq libraries from the remaining 260 human DNA samples, which were subjected to deep and shallow sequencing. With the TIME-Seq Human Blood Clock, we observed similar predictive accuracy in the validation dataset compared to the initial testing set (*R*=0.96; Fig. 6h). Using the initial data for reference, *scAge* predictions were also highly accurate (*R*=0.93) when applied to validation shallow sequencing data (median N=39,846 reads) using up to the 10^th^ percentile of top age-correlated CpGs (Fig. 6i-j). Age-adjusted residuals for the TIME-Seq Human Blood Clock and *scAge* predictions were significantly correlated (*R*=0.52, p<2.2e-16), suggesting the methods reflect a common underlying biological signal.

## Discussion

Epigenetic clocks are increasingly ubiquitous tools for both clinical and basic aging research. Until now, they have been relatively expensive and laborious to measure, limiting their application to modest-sized or extremely well-funded experiments. TIME-Seq is a flexible and scalable targeted sequencing approach that decreases costs of epigenetic clocks by up to 100-fold in large experiments. Using TIME-Seq, we built and validated epigenetic clocks to predict age in over 2700 unique samples. This scale was enabled by immediate sample barcoding that facilitates low-cost, pooled library preparation compatible with efficient hybridization-based enrichment and bisulfite conversion. Compared to traditional RRBS library preparations or BeadChip sample preparation, which can take anywhere from 4-9 days^43^ and cost upwards of $30-$50 per sample, reagent cost for TIME-Seq libraries is only $0.65 per sample, and without any automation, a trained technician can easily prepare upwards of 800 samples in only 1.5 days (approximately 12 hours of hands-on time). Further, input DNA for TIME-Seq (100 ng) is the same as standard RRBS libraries and 3-5 times less than DNAme microarray, enabling longitudinal measurement of epigenetic age from low-yield DNA extractions such as a mouse cheek bleed.

Using a combination of shallow-TIME-Seq and *scAge*, we show that it is possible to accurately predict age in bulk samples from around ten thousand sequencing reads per sample. We applied shallow-TIME-Seq with *scAge* to predict age in mouse and human blood and mouse liver, finding it surprisingly accurate with correlations nearing those of elastic-net based clocks. Even in smaller scale experiments, using *scAge* in combination with TIME-Seq library preparation and shallow sequencing, the cost of accurate age prediction was more than 100-fold lower than conventional methods. While *scAge* predictions were slightly less robust than clock predictions, future optimizations will likely increase accuracy and allow for ultra-cheap age prediction in extremely large studies.

While highly targeted approaches have been described for epigenetic age prediction^44-46^, such as pyrosequencing or digital PCR, these methods are more expensive and less scalable than TIME-Seq-based predictions from our clocks or *scAge* predictions from shallow TIME-Seq. Further, their reliance on low-CpG clocks (e.g., 3-15 CpGs) limits the number of aging modules they reflect. TIME-Seq is flexible and capable of high enrichment and deep coverage of thousands to tens-of-thousands of CpGs. Further, the cost for accurate TIME-Seq clocks gets substantially lower at larger scales. Conservatively, we estimate that 12,500 samples could be prepared and sequenced for clock prediction on a single NovaSeq S4 flow cell for just $1.70 per sample. With shallow sequencing of TIME-Seq libraries followed by *scAge*, per-sample costs for age prediction drop to just 87¢.

A central goal of aging research has been to develop biomarkers capable of detecting differences in the biological rate of aging^1^. We show that TIME-Seq clocks can be used to detect the effect of interventions that accelerate, slow, or reverse the pace of aging *in vitro* and *in vivo*. Designed for low-cost and scale, TIME-Seq is uniquely capable of biomarker application to large intervention studies such as screens or longitudinal clinical trials. Whether or not TIME-Seq clocks or *scAge* will detect age-altering effects in every tissue, or if they will associate with other metrics of functional decline, will require additional experiments. For instance, our analysis of TIME-Seq clocks in mouse blood revealed the lack of correlation between epigenetic age and frailty described in human clocks^34^, suggesting instead that DNA methylation biomarkers might be specifically designed to predict frailty-adjusted age in mice.

In the current study, we rationally designed hybridization probes to enrich for loci that were already known to correlate with chronological age, our target metric. However, when such data is not available, more genomic area might be enriched to identify CpGs with high-correlation relative to the phenotype of interest (e.g., mortality, frailty, or cancer status). These libraries would initially be more expensive to sequence, but the number of baits could be reduced once model CpGs were discovered. This separation of more expensive biomarker discovery from low-cost measurement will be key to the widespread adoption and routine use of DNAme clocks as well as clocks based on other biomolecules^7,47^.

Being designed for minimal cost and maximal efficiency, TIME-Seq has the same trade-offs as other Tn5-based library preparations. For example, samples with improperly normalized DNA tend to drop-out due to either over-or under-tagmentation. During this study, we developed increasingly automated and reliable approaches for DNA normalization at scale (see Methods), which limited sample drop out. Sample normalization could also be addressed using on-bead tagmentation^48^ to control for variation in starting DNA content, but this would require modifications to the adaptor design. Ultimately, further automation of our TIME-Seq sample processing pipeline could enable a single scientist to prepare thousands of samples at once, enabling low-cost age predictions from extremely large cohorts such as the U.K. Biobank. Such a study would be a powerful resource to identify genetic and lifestyle factors that influence aging at population scale.

## Methods

### DNA extraction, quantification, and normalization

For blood, 100-300 μl of pelleted mouse whole blood (plasma removed) was resuspended in 1mL of red blood cell (RBC) lysis buffer (155 mM NH4Cl, 12 mM NaHCO3, 0.1 mM EDTA, pH 7.3), incubated for 10 minutes on ice, and centrifuged at 2000 RCF for 5 minutes. Pelleted cells were resuspended in 1.5mLs of RBC lysis buffer and spun twice before being lysed in 800 μl TER (50 mM Tris-HCl pH8, 10 mM EDTA, 40 μg/mL RNase A) with 50 μl of 10% SDS added. Lysates were incubated at 37ºC to allow for RNA degradation and then 50 μl of 20 mg/mL proteinase K was added and samples were incubated overnight at 65ºC. To purify DNA, 500 μl of 1:5 diluted (dilution buffer: 20% PEG 8000, 2.5 M NaCl, 10 mM Tris-HCl pH8, 1 mM EDTA, 0.05% Tween-20) SPRI DNA binding beads were added to each sample, and they were incubated with rotation for 30 minutes at room temperature. Tubes were then placed on magnetic racks to capture SPRI beads, and the beads were washed with 1 mL of ice-cold 80% ethanol twice. DNA was eluted in 75 μl of 10mM Tris-HCl (pH 8). Purified DNA was quantified using the Qubit double-stranded DNA broad range kit (Catalog No. Q32850, ThermoFisher) and diluted to 10 ng/μl for TIME-Seq reactions. To check for contaminants, a subset of samples from each extraction were assessed by NanoDrop. For tissues, frozen tissues from mice of various ages were collected from our internal mouse aging tissue bank. Tissues were kept frozen on dry ice while 10-20mg of tissue was cut into a well of a 96-well plate. DNA was extracted using the Chemagic 360 system (Perkin Elmer) and the 10 mg tissue kit (Perkin Elmer, CMG-723) and resuspended in 100 μL of elution buffer.

To achieve consistent DNA normalization from four 96-well DNA plates at a time, we developed a scalable and partially automated protocol to get concentrations of 10 ng/μL with a low-percentage drop-out due to over-or under-dilution. To begin, a standard starting volume of approximately 25 μL of DNA was transferred to a 96-well plate (BioRad, HSP9601). 1μL of unnormalized DNA was removed and transferred to a black 96-well plate (VWR, 33501-812) with 99μL of TE (10mM Tris, 1mM EDTA). 100μL of 1X picogreen solution (ThermoFisher, P7589) was added to the DNA, after which the plates were sealed, vortexed for 30 seconds, and spun in a plate spinner. Fluorescence was measured by the SpectraMax i3 (Molecular Devices). Standards in triplicate from 1-100ng of total DNA were used to predict the amount of DNA from each sample. Using these DNA concentrations, samples were diluted with 10 mM Tris pH 8 using the Mantis liquid handler (Formulatrix) to a concentration of 40 ng/μL if ≥ 100 ng/μL or 20 ng/μL if < 100 ng/μL. After a second round of DNA quantification, samples were then transferred to a new 96-well plate at a standard volume (typically 20-30 μL) using a BioMek FXp liquid handler (Beckman Coulter). This plate was diluted to a final TIME-Seq starting concentration of 10 ng/μL using the Mantis liquid handler.

### Tn5 transposase purification

Tn5 transposase was purified according to a described optimized protocol^49^. Briefly, 1 mL of overnight culture containing pTXB1-Tn5 (Addgene plasmid #60240) was used to inoculate 1 L of ZYM-505 growth media containing 100 μg/mL ampicillin and 0.001% polypropylene glycol (L14699-AE, Alfa Aesar). After the culture was grown for 4 hours at 37ºC, IPTG (0487-10G, VWR) was added to 0.25 mM, and the culture was grown for an additional 4 hours at 18ºC. The culture was centrifuged at 25,000 RPM for 25 minutes, and the pelleted culture was flash-frozen in liquid nitrogen before being stored overnight at −80ºC. The pellet was thawed on ice, resuspended in 10 mL of HEGX buffer (20mM HEPES-KOH pH 7.2, 0.8 M NaCl, 1mM EDTA, 10% glycerol, 0.2% Triton X-100) with a Roche protease inhibitor (SKU11697498001, Millipore Sigma), and 1% w/v of pre-dissolved (50% w/v) octyl-thioglucoside (O-130-5 Gold Bio) was added to help lysis. After a 10-minute incubation on ice, 100 mL of HEG-X was added, and the lysate was transferred to a glass beaker for sonication on the 550 Sonic Dismembrator (ThermoFisher) using 15 cycles (15 seconds on, 30 seconds off) on 70% duty and power 7. The sonicated lysate was pelleted at 30,000 RPM for 30 minutes at 4ºC. The supernatant was transferred to a clean beaker with a stir bar, placed on a magnetic stir plate, and 10% PEI was added dropwise while stirring to remove excess bacterial DNA. After 15 minutes, PEI and precipitated DNA were removed by spinning the mixture at 30,000 RPM for another 30 minutes at 4ºC. The lysate was added to 2 chromatography columns (7321010, BioRad) packed with 25 mL each of chitin resin (S6651S, NEB) and equilibrated with 100 mL of HEG-X each. The supernatant was added to the columns in equal proportion and allowed to flow through, before the column was washed with 30 column volumes of HEG-X. To elute the purified Tn5, 25 mLs of HEG-X with 100 mM DTT was added to each column and 10 mLs was allowed to flow through before sealing the column and letting it stand for 44 hours. 27 mL of elution was collected, and a Bradford assay (23200, ThermoFisher) was used to quantify protein. The elution was then concentrated to 20 mL using Amplicon Ultra 30 kDA filters (UFC900308, Millipore Sigma) and dialyzed in a Slide-A-Lyzer (66212, ThermoFisher) cassette with 1 L dialysis buffer (50 mM HEPES-KOH, pH 7.2 0.8M NaCl, 0.2 mM EDTA, 2 mM DTT, 20% glycerol). After two rounds of dialysis totaling 24 hours, the eluted protein was removed, aliquoted into 1 mL microcentrifuge tubes, and flash frozen in liquid nitrogen before being stored at −80ºC. The final concentration of purified protein was 1.5 mg/mL, and we estimate that 1L of culture produced enough Tn5 for approximately 16,000 TIME-Seq reactions with 100 ng of DNA per sample.

### Activation of TIME-Seq Tn5

Oligonucleotides (oligos) were ordered from IDT and HPLC purified except for TIME-Seq indexed adaptors and hybridization blocking oligos (see Supplementary Table 5 for all oligonucleotide sequences). To anneal TIME-Seq adaptors, 100 μM TIME-Seq adaptor B containing a 5-bp internal barcode and (separately) 100 μM methylated adaptor A were combined in equal volume with the 100 μM Tn5 reverse ME oligo. Enough methylated A adaptor was annealed to be added in equal proportion with each indexed adaptor B. Oligos were denatured at 95ºC for 2 minutes and then ramped down to 25ºC at 0.1ºC/s. The annealed oligos were then diluted with 85% glycerol to 20 μM, and the methylated A adaptor, indexed TIME-Seq B adaptor, and 50% glycerol were combined in a ratio of 1:1:2. The resulting 10μM adaptors were combined in equal volume with purified Tn5 (1.5 mg/mL), mixed thoroughly by pipetting 20 times, and incubated at room temperature for 50 minutes. Activated transposomes were stored at −20ºC and no loss of activity has been observed up to 8 months.

To test activity of TIME-Seq transposomes, 100 ng of human genomic DNA (11691112001, Roche) was tagmented in 25 μl reactions by adding 12.5 μl of 2X tagmentation buffer (20mM Tris-HCl pH 7.8, 10 mM dimethylformamide, 10 mM MgCl_2_) using 1.5 μl of each barcoded transposome. After reactions were incubated at 55ºC for 15 minutes, STOP buffer (100mM MES pH5, 4.125M guanidine thiocyanate 25% isopropanol, 10mM EDTA) was added to denature Tn5 and release DNA. To assess tagmentation, 90% of each reaction was run in separate lanes of a 1% agarose gel at 90 volts (V) for 1 hour. The gel was stained with 1x SYBR gold (S11494, ThermoFisher) in Tris-Acetate-EDTA (TAE) buffer, and DNA fragment size was determined using a ChemiDoc (Bio-Rad) for gel imaging. The remaining DNA was pooled, cleaned up using a DNA Clean & Concentrator-5 (D4013, Zymo) kit, and the DNA was amplified with barcoded TIME-Seq PCR primers. Amplified pools were spiked into sequencing runs to assess relative barcode activity when new transposase was activated.

### Biotin-RNA bait design and production

Mouse and human discovery baits were designed to enrich for previously described epigenetic clock CpGs^11-13,50^ using mm10 and hg19 genomic coordinates. Using bedtools^51^ (v2.28.0), regions around each target CpG were first expanded 125 base pairs (bps) up-and downstream (*bedtools slop*) and overlapping loci were merged (*bedtools merge*). Next, 110 bp bait windows were defined every 20 bps in the region (*bedtools window*) and the baits were intersected (*bedtools intersect*) with a file containing RepeatMasker (http://www.repeatmasker.org) annotated regions to identify baits overlapping repetitive DNA regions. Next, the FASTA sequence of each bait was gathered (*bedtools getfasta*), and blat (version 35) was used to get the copy number for each bait with options, -fastMap -maxIntron=50 -stepSize=5 -repMatch=2253 - minScore=40 -minIdentity=0. Using custom R (v. 4.0.2) scripts, information was gathered from the output files from each probe including the percent of each nucleotide, the overlap with repeats, and the bait copy number as determined by blat. Baits were automatically filtered that had overlap with repeats, however, each locus that had none or very few (< 4) baits after filtering were inspected manually and, if the blat copy number was low (< 10), baits were added back, either to the exact locus or shifted slightly to avoid the annotated repeat. In preliminary biotin-RNA hybridization experiments (data not shown), we noticed that baits with high or low percentage T (less than 8% or greater than 30%) had low coverage, possibly due to stalling of the RNA polymerase while incorporating biotin-UTP. Therefore, when more than half the baits at a target locus had a percent T greater than 30% or lower than 8%, the reverse complement strand was captured instead for all baits at the locus.

To design enrichment baits for mouse and human rDNA, FASTA sequences were prepared from GeneBank accessions BK000964.3 (mouse) and U13369.1 (human) according to previously described methods^28^ by moving the last 500 bps of each sequence (rDNA promoter) to be in front of the 5’ external transcribed spacer (ETS). From the region comprising the promoter to the 3’ ETS (mouse, 13,850bps; human, 13,814bps), 110-nt baits were designed using the *bedtools window* function to create baits every 20bps. Version 1 rDNA baits used in the pilot and targeting the original rDNA clock were designed to specifically enrich rDNA clock CpGs^28^ using the same approach described for non-repetitive clock CpGs (i.e., 250bp windows were merged and 110-nt baits were designed to tile the regions).

The sequence of each bait set was appended with a promoter (Sp6 or T7) for *in vitro* transcription (IVT), as well as a promoter-specific reverse priming sequence that contained a BsrDI restriction enzyme motif. Bait sets containing Sp6 and T7 promoters were ordered together in a single-stranded DNA oligo pool (Twist), and pools were resuspended to 10 ng/μl. Bait sets were amplified in reactions containing 12.5 μl of 2X KAPA HiFi HotStart Polymerase Mix (7958927001, Roche), 0.75 μl of each 10 μM primer, and 0.5 μl of the bait pool using the following thermocycler program: initial denaturation, 95ºC for 3 min; 10 cycles amplification, 98ºC for 20 seconds, 61ºC (Sp6) or 58ºC (T7) for 15 seconds, 72ºC for 15 seconds; a final elongation for 1 minute at 72ºC. Amplified DNA was cleaned up using a Clean & Concentrator-5 kit (D4013, Zymo) and then digested with 1 μl BsrDI (R0574S, NEB) at 65ºC for 30 minutes. This reaction was again purified with a Clean & Concentrator-5 kit, and IVT reactions were set up according to the HiScribe™ T7 (E2040S, NEB) or Sp6 (E2070S, NEB) High Yield kits using half of the cleaned DNA template for each reaction and storing the rest at −80ºC. All ribonucleotides were added to a final concentration of 5mM, including a 1:4 ratio of biotin-16-UTP (BU6105H, Lucigen) to UTP at 5 mM. After the reactions were incubated for 16 hours overnight at 37ºC, 25 μl of nuclease free water and 2 μl of DnaseI (M0303S, NEB) were added to degrade the DNA template. RNA was purified using the 50 μg Monarch® RNA Cleanup Kit (T2040S, NEB). The concentration of RNA was measured using Qubit RNA BR Assay Kit (Q10210, ThermoFisher) and the size of RNA was measured using an RNA ScreenTape (5067-5576, Agilent) on an Agilent Tapestation.

While the yield from each DNA amplification and IVT reaction varies depending on the size and composition of the bait sets, we estimate that 1,000-10,000 hybridization reactions-worth of bait could be produced from just 1 single-stranded oligo pool. For TIME-Seq libraries of 48-64 samples, this could enrich tens to hundreds of thousands of samples. Another advantage of this approach is that baits can be easily shared with other researchers as the ssDNA template, amplified DNA, or prepared biotin-RNA.

### TIME-Seq library preparation

For TIME-Seq library preparations, samples were organized into relatively even pools, and 10 μl of DNA (10 ng/μl, 100 ng total) from each sample was distributed into separate wells of strip tubes (or 96-well plates) for tagmentation. 100 ng of unmethylated lambda phage DNA (D1521, Promega) was tagmented with each pool. Lambda DNA that came through at a low percentage of demultiplexed reads served to estimate bisulfite conversion efficiency. To tagment samples, 12.5 μl of 2X tagmentation buffer (20mM Tris-HCl pH 7.8, 10mM dimethylformamide, 10mM MgCl_2_) was added to each sample. Next, 2.5 μl of uniquely indexed TIME-Seq transposome was added, and the reaction was immediately mixed by pipetting 20 times. Once transposomes were added to each sample in a pool, the samples were placed at 55ºC for 15 minutes. After incubation, 7 μl of STOP buffer (100mM MES pH5, 4.125M guanidine thiocyanate 25% isopropanol, 10mM EDTA) was added, pools were vortexed and pulse spun in a centrifuge, and the reaction was incubated at 55ºC for an additional 5 minutes.

After stopping the reactions, samples from each pool were combined into a single tube, typically a 5 mL Lo-bind tube (0030122348, Eppendorf) or 15 mL falcon tube (229410, Celltreat), and 118 μl per sample of DNA Binding Buffer (D4004-1-L, Zymo) was added. Pools were then applied to Clean & Concentrate-25 (D4033, Zymo) columns. If the volume of the pool exceeded 5 mL, each pool was passed in equal volume through 2 separate columns. After 2 washes, pools were eluted in 41 μl (typically yielding 39 μl after elution) and 1 μl was removed to assess tagmentation fragment size and yield by D5000 ScreenTape (Catalog No. 5067-5588, Agilent) on an Agilent Tapestation.

For methylated end-repair, eluted pools were combined with 5 μl of NEB Buffer 2, 5 μl of 5mM dNTPs containing 5-methyl-dCTP (N0356S, NEB) instead of dCTP, and 2 μl of Klenow Fragment (3’→5’ exo-) (M0212L, NEB). The reactions were incubated at 37ºC for 30 minutes and then cleaned up with a Clean and Concentrate-25 column (Zymo). To elute pools, 30 μl of heated elution buffer was applied to the column and incubated for 1 minute before being spun. Eluted DNA was then passed through the column a second time and concentrated for hybridization to 5 μl with a SpeedVac Concentrator (Eppendorf).

For each pool, DNA, RNA, and hybridization mixtures were prepared in separate strip tubes (1 per pool). On ice, DNA mixtures were prepared by adding 5 μl of concentrated tagmented DNA from each pool, 3.4 μl of 1μg/μl mouse cot-1 (18440016, ThermoFisher) or human cot-1 (15279011, ThermoFisher), and 0.6 μl of 100 μM TIME-Seq hybridization blocking primers (IDT). RNA mixtures were prepared on ice by combining 4.25 μl of nuclease free H2O with 1 μl of Superase•In RNase inhibitor (AM2696, ThermoFisher), mixing, and then adding 0.75 μl (750 ng total) of the biotin-RNA baits. Hybridization mixtures were kept at room temperature and comprised 25 μl 20X SSPE (AM9767, ThermoFisher), 1 μl 0.5 M EDTA, 10 μl 50X Denhardt’s Buffer (1% w/v Ficoll 400, 1% w/v polyvinylpyrrolidone, 1% w/v bovine serum albumin), and 13 μl of 1% SDS. Once the mixtures were prepared for each pool, the DNA mixtures were placed in a thermocycler and incubated for 5 minutes at 95ºC. Next, the thermocycler cooled to 65ºC, and the hybridization mix was added to the thermocycler. After 3 minutes at 65ºC, the RNA mix was added to the thermocycler and incubated for 2 minutes at 65ºC. Next, the thermocycler lid was opened and, keeping all tubes in the thermocycler well, 13 μl of heated hybridization buffer was transferred to the RNA baits mixture, followed by 9 μl of the denatured TIME-Seq pooled DNA. This step was done quickly to limit temperature change during transfer, typically with a multichannel pipette for multiple pools. The combined mixtures were pipetted to mix 3-5 times, capped, and the thermocycler lid was closed. The hybridization reaction was then incubated at 65ºC for 4 hours.

To capture biotin-RNA:DNA hybrids, 125 μl of streptavidin magnetic beads were washed three times in 200 μl of binding buffer (1M NaCl, 10mM Tris-HCl pH 7.5, 1 mM EDTA) and resuspended in 200 μl of binding buffer. With the reaction still in the thermocycler, the streptavidin beads were added to the reactions and then quickly removed to room temperature. The reactions were rotated at 40 RPM for 30 minutes to allow for biotin-streptavidin binding and then placed on a magnetic separation rack (20-400, Sigma-Aldrich) until the solution was clear. Next, the beads were resuspended in 500 μl of hybridization wash buffer 1 (1X SSC, 0.1% SDS) and incubated at room temperature for 15 minutes. The beads were separated again on the magnetic separation rack and quickly resuspended in 500 μl of pre-heated 65ºC wash buffer 2 (0.1X SSC, 0.1% SDS), then incubated for 10 minutes at 65ºC. This step was repeated for a total of 3 heated washes. On the final wash, beads were magnetically separated, resuspended in 22 μl of 0.1M NaOH, moved to a new strip tube (to avoid droplets on the original tube mixing with the elution buffer), and incubated for 10 minutes. After 10 minutes, beads were separated and 21 μl of the eluted ssDNA from each pool was moved to another new strip tube for bisulfite conversion reaction.

Bisulfite conversion was done using the EpiTect Fast Bisulfite Conversion Kit (59824, Qiagen), which was chosen due to the inclusion of DNA protect buffer (limiting DNA strand breakage in bisulfite reaction) and carrier RNA that helps yield from low-concentration reactions. The volume of the eluted DNA was adjusted to 40 μl using nuclease free water, 85 μl of Bisulfite Solution was added, followed by 15 μl of the DNA Protect Buffer, and the solution was mixed thoroughly. Bisulfite conversion and clean-up proceeded according to standard kit instruction. DNA was eluted in 23 μl of kit elution buffer. The initial elution was passed through the column a second time.

PCR amplification was done in a 50 μl reaction containing 23 μl of the eluted DNA, 1 μl of 25 μM P7 indexed primer (see Supplementary Table 5 for primer sequences), 1 μl of 25μM P5 indexed primer, and 25 μl NEB Q5U 2X Master Mix. Reactions were amplified with the following program: initial denaturation at 98ºC for 30 second; 19 cycles of 98ºC for 30 seconds, 65ºC for 30 seconds, and 72ºC for 1 minute; a final elongation at 72ºC for 3 minutes. After PCR reactions were finished, they were cleaned using 1.8X CleanNGS SPRI Beads (CNGS005, Bulldog-Bio). Library fragment size and yield was assessed using a D1000 (5067-5582, Agilent) or High Sensitivity D1000 ScreenTape (5067-5584, Agilent) on an Agilent Tapestation. Pools were combined for sequencing.

### Sequencing

TIME-Seq library sequencing requires two custom sequencing primers (see Supplementary Table 5) for read 2 (Tn5 index) and index read 1 (i7 index), which were spiked into standard primers for all sequencing runs so that standard or control libraries (e.g., PhiX) could be readout as well. A complete list of sequencing platforms used for each experiment is contained in Supplementary Table 3. TIME-Seq libraries that were sequenced on an Illumina MiSeq using a 150 cycle MiSeq v3 kit (MS-102-3001, Illumina) or on a NextSeq 500 using a 150 cycle NextSeq High (20024907, Illumina) or Mid (20024904, Illumina) Output v2.5 kit used the following read protocol: read 1, 145-153 cycles; i7 index read, 8 cycles; i5 index read (if needed), 8 cycles; read 2, 5 cycles. This custom read protocol helps maximize sequencing efficiency and decrease cost, since the fragment size of amplified TIME-Seq libraries were typically 80-200bps and sequencing with larger kits results in a large portion of overlapping (unused) cycles. Optimization experiments, smaller deep-sequenced pools, and shallow sequencing data were typically generated by sequencing on a MiSeq using a MiSeq Reagent Micro v2 kit (MS-103-1002, Illumina) using standard paired-end and dual indexed read cycle numbers. Larger experiments sequenced on a NovaSeq 6000 using a 200 cycles NovaSeq 6000 SP or S1 Reagent Kit v1.5. High GC genome from *Deinococcus radiodurans* was spiked into sequencing runs in different proportions (1-3% on MiSeq, 5-10% on NovaSeq, and 15-20% on NextSeq) to increase base diversity and improve sequencing quality^52^.

### Sequenced read demultiplexing and processing

TIME-Seq pools were demultiplexed using sabre (https://github.com/najoshi/sabre) to identify the internal Tn5 barcode with no allowed mismatches and separate reads into unique FASTQ files for each sample. Cutadapt (version 2.5) was used to trim adaptors (PE: -G AGATGTGTATAAGAGANAG -a CTNTCTCTTATACACATCT -A CTNTCTCTTATACACATCT; SE option: -a CTNTCTCTTATACACATCT). Reads were mapped using Bismark^53^ (version v0.22.3; options -N 1 --non_directional) to bisulfite converted genomes (*bismark_genome_prepararation*) mm10, hg19, or custom rDNA loci (see bait design methods), and reads were subsequently filtered using the bismark function *filter_non_conversion* (option --threshold 11). Importantly, the latter step does not (as the Bismark function name suggest) reflect non-conversion from sodium bisulfite treatment, rather it removes a small percentage of reads (0.5%-3%) that are artificially fully methylated during the methylated end-repair of the reads, which has been previously described^52^. Next, the bismark function *bismark_methylation_extractor* was used to call methylation for each sample with options to avoid overlapping reads (--no-overlap) in PE sequencing and to ignore the first 10 bps of each read (if SE: --ignore 10 and --ignore_3prime 10; if PE: --ignore_r2 10 --ignore_3prime_r2 10 as well), which precludes bias from methylated cytosines added in the Tn5 insertion gap during end-repair. Bisulfite conversion efficiency was assessed from unmethylated lambda DNA mapped to the bisulfite-converted *Enterobacterphage lambda* genome (iGenomes, Ilumina) and was generally ≥99%.

### Epigenetic clock training, testing, and analysis in validation and intervention cohort data

R (version 4.0.2) was used for all data analysis, including data organization, clock training and testing, and applying clocks to validation data. For clock training, methylation and coverage data were taken from bismark.cov files (output of *bismark_methylation_extractor*), which contain CpG location and methylation percent (0-100). Since high-coverage rDNA loci have been shown to make better age prediction models^21^, mouse rDNA methylation data was filtered to comprise only CpGs with high coverage (≥ 200) in greater than 90% of samples at each CpG in the coverage matrix. For other mouse and human clock CpG enrichment datasets, CpGs were filtered to have at least coverage 10 in 90% of the samples.

To build epigenetic clocks from deep-sequenced TIME-Seq data, samples were randomly selected from discrete age groups (e.g., 25-55 weeks old for mice, 30-40 years old for humans) in approximately 80:20 training to testing ratio. From the training set data, penalized regression models were trained to predict age from methylation values with the R package glmnet^54^ with alpha set to 0.05 or 0.1 (elastic net). To further refine the model, age predictions from the training data were regressed against age and the coefficients from this simple linear model were included to adjust the elastic net model (see full equation below for computing clock predictions). While minor, this adjustment helped to correct for over- or under-prediction in the youngest and oldest samples and produced predictions with more normal ΔAge across lifespan. Multiple elastic net models were trained using random train-test splits, and a model with high Pearson correlation and low median error in the testing set was selected for application to the independent validation cohorts or intervention samples. This same process was applied to build the RRBS-TS-rDNA clock, filtering for CpGs in TIME-Seq data that had minimum coverage of 50 in the RRBS mouse blood clock dataset^11^.

Clocks were applied to validation and intervention samples by joining bismark.cov files with the corresponding clock CpG coefficients. When a clock CpG was not covered, the missing value was replaced by the average methylation percent at that CpG from all other samples in the experiment. Samples were excluded if >10% of clock CpGs were missing, or samples contained less than 100,000 reads. To calculate the weighted methylation (*S*) from elastic net regression coefficients, each clock CpG methylation percent is multiplied by the corresponding clock coefficient and these values are summed, as follows:

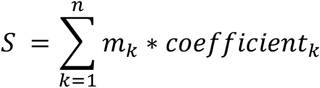

Where *n* is the total number of clock CpGs, and *m*_*k*_ is a number from 0-100 representing percent methylation. To calculate predicted age, the intercept from elastic net regression is added to *S*, and this value is adjusted with the simple linear regression coefficients *a* and *c*, as follows:

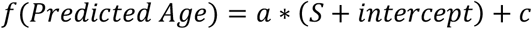

Clocks predict age in units of weeks for the TIME-Seq Mouse rDNA Clock, months for all other mouse clocks, and years for the TIME-Seq Human Blood Clock. See Supplementary Table 6 for clock positions, coefficients, and intercepts.

### *scAge*-based epigenetic age predictions from shallow TIME-Seq

To obtain accurate epigenetic age predictions from shallow TIME-seq data, we employed the use of the recently introduced *scAge* framework^25^. Unlike to the original *scAge* application in single-cell data, all methylation values from shallow sequencing were retained unmodified (i.e., without introducing a forced binarization step for fractional methylation values). We used bulk blood- and liver-specific methylation matrices to compute linear regression equations and Pearson correlations between methylation level and age for each CpG within a particular training set. These equations were in the form:

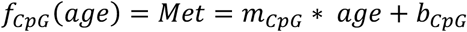

where *age* is treated as the independent variable predicting methylation, and *m* and *b* are the slope and intercept of the CpG-specific regression line, respectively. This enabled the creation of reference linear association metrics between methylation level and age for each CpG covered in the training datasets. Five separate training datasets were used to compute epigenetic ages in shallow sequencing data: (1) Bulk blood RRBS methylation profiles from 50 normally-fed C57BL/6J mice from the Thompson et al. (2018) study across 1,202,751 CpG sites; (2) Bulk liver RRBS methylation profiles from 29 normally-fed C57BL/6J mice from the Thompson et al. (2018) study across 1,042,996 CpG sites; (3) Bulk blood RRBS methylation profiles from 153 normally-fed C57BL/6J mice from the Petkovich et al. (2017) study across 1,918,766 CpG sites; (4) Bulk blood TIME-seq methylation profiles from 120 normally-fed C57BL/6J mice from the present study across 6,884 CpG sites; (5) Bulk liver TIME-seq methylation profiles from 103 normally-fed C57BL/6J mice from the present study across 12,480 CpG sites; and (6) Bulk human blood TIME-Seq methylation profiles from 796 samples across 9379 CpGs. Methylation values from the Thompson et al. (2018) study were concatenated to the positive strand, as described in the original study.

We intersected individual methylation profiles of shallow TIME-seq data with the reference datasets, producing a set of *n* common CpGs shared across both datasets. For each shallow sample, we filtered these *n* CpGs based on the absolute value of their correlation with age (|*r*|) in the reference data, selecting the most age associated CpGs in every sample. We opted to use a percentile-based approach to select informative CpGs for predictions, which takes into account differential coverage across samples. Furthermore, this enabled more consistent correlation distributions among shallow TIME-Seq profiles of different coverage.

For each selected CpG per sample, we iterated through age in steps of 0.1 months for mice or 0.1 years for humans from a minimum age to a maximum age value. We iterated from -20 months to 60 months for mice and -50 years to 150 years for humans to encompass a wide range of possible predictions without artificially bounding outputted predicted values. We observed in our testing that all predictions generated by *scAge* for these data were within this generous range and not near the extremes. Using the linear regression formula calculated per individual CpG in a reference set, we computed the age-dependent predicted methylation, *f*_*CpG*_(*age*), which by the nature of the data normally lies between 0 or 1. If this predicted value was outside of the range (0, 1), it was instead replaced by 0.01 or 0.99 depending on the proximity to either value. This ensured that predicted bulk methylation values were bounded in the unit interval, corresponding to a biologically meaningful range between fully unmethylated and fully methylated.

Next, we computed the probability of observing a particular methylation state at every age in the given range based on the reference linear model estimate. For this, we calculated the absolute value of the distance between the observed methylation fraction in the shallow sample and the estimated methylation value from the reference linear model. Next, we subtracted this absolute distance from 1; hence, the closer the observed value is from that predicted by the linear model, the higher the probability of observing this state at a particular age. This provided an age-dependent probability for every common CpG retained in the algorithm.

Lastly, assuming that all CpGs in a particular sample are independent from each other, the product of each of these CpG-specific probabilities will be the overall probability of the observed methylation pattern: 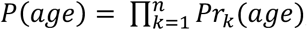 where *k* represents individual CpGs. We then found the maximum of that product among different ages (i.e., to find the most probable age for observing that particular methylation pattern). To do this, we compute the sum across CpGs of the natural logarithm of the individual age-dependent probabilities, preventing underflow errors when many CpGs are considered. This gave us 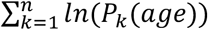 for each age step. By harnessing the relationship of methylation level and age at many CpGs, these logarithmic sums provide a single likelihood metric for every age step within the defined bounds. We picked the age of maximum likelihood as our predictor of epigenetic age for a particular shallow TIME-Seq sample.

### Comparison between age prediction methods in the same mouse or sample

While age predictions were highly accurate, we still observed slight bias in prediction for the oldest and youngest samples, which could influence the correlation between predictions made in the same mouse or sample. To control for this bias and compare predictions, we regressed predicted ages against chronological age for the entire set of data (e.g., the testing set from clock training, or the entire validation set) using the *lm()* function in R and calculated the residual for each prediction. For brevity, we refer to these values as the age-adjusted prediction residuals in our Results section.

### Animal assessments

All mouse experiments were approved by the Institutional Animal Care and Use Committee of the Harvard Medical Area Standing Committee on Animals. Male and female C57BL/6Nia mice were obtained from the National Institute on Aging (NIA, Bethesda, MD), and group housed (3-4 mice per cage) at Harvard Medical School in ventilated microisolator cages with a 12:12 hour light cycle, at 71°F with 45-50% humidity. Mouse blood samples (150-300 μl) were collected in anesthetized mice (3% isoflurane) from the submandibular vein into tubes containing approximately 10% by-volume of 0.5M EDTA. Blood was spun at 1500 RCF for 10 minutes and plasma removed. Blood cell pellets were stored frozen at − 80ºC. For validation experiments, a sub-sample of whole blood was stored on ice and processed within 4 hours with the Hemavet 950 (Drew Scientific) to give 20 whole blood count parameters. Frailty was assessed using the mouse clinical frailty index^31^, a non-invasive assessment of 31 health deficits in mice. For each item, mice were scored 0 for no deficit, 0.5 for a mild deficit, and 1 for a severe deficit. Scores were added and divided by 31 to give a frailty index between 0 and 1, where a higher score indicates a greater degree of frailty. For more details see http://frailtyclocks.sinclairlab.org. 200 mouse (C57BL/6N) ocular-vein blood samples were collected by researchers at The Jackson Laboratory’s Nathan Shock Center (Bar Harbor, Maine) according to methods described previously^55^. These samples were used for TIME-Seq clock training and testing, as well as shallow-sequencing analysis.

### Benchmarking TIME-seq against BeadChip and RRBS

For benchmarking experiments, 48 mouse blood DNA samples (independent from the validation set) were prepared with TIME-Seq using mouse clock enrichment baits. 500 ng of DNA from the same 48 samples was sent to FOXO Technologies for analysis on the Infinium Mouse Methylation BeadChip (Illumina, 20041559). 18 of these DNA samples with at least 100 ng of DNA remaining were prepared and analyzed with RRBS using published library preparation methods^12^. RRBS samples were pooled and sequenced on an Illumina NovaSeq to a median depth of 38 million read pairs and mapped with Bismark (version v0.22.3) and methylation data was extracted by *bismark_methylation_extractor* with option --no_overlap.

For analysis of CpG methylation levels, loci common to each sample from each method (and >20 coverage for sequencing approaches) were identified, and the methylation levels from each technology were plotted pairwise as shown in Figure 4. For clock analysis, the TIME-Seq mouse blood and multi-tissue clocks were applied to TIME-Seq data as described above, whereas the RRBS-based mouse blood clock was applied in a similar fashion using the reported loci, weights, and formula^12^, replacing any missing values with the mean methylation from all other samples. The BeadChip clock was applied to BeadChip data as reported^56^.

For comparison of costs at various scales, consumable costs were estimated (without consideration of labor) as follows: (1) TIME-Seq costs were estimated as the library preparation cost per sample ($0.65, detailed in Supplementary Table 1) and the cost of sequencing reagents sufficient to sequence each sample to at least 750,000 reads, which is 50% more than the minimum depth we report for accurate age estimation. (2) For BeadChip, costs were estimated as $25 per sample for DNA preparation below 1000 samples and $20, $17.5, and $15 per sample for volumes of 4992, 9984, and 12,000 samples. The cost of consumables for BeadChip assay were taken from the list price for Infinium Mouse Methylation BeadChip on Illumina’s website (Illumina, catalog numbers 20041558, 2004159, 20041560). (3) For RRBS, sample preparation costs were estimated as $50 per sample below 1000 samples and $30 per sample for volumes of 1000 and 5000 samples, and $20 per sample for volumes of 10,000 and 12,500 samples. Costs of sequencing reagents were estimated to provide median reads per sample of 45 million.

### Mouse intervention studies

For the late-life dietary interventions, male and female C57BL/6Nia mice were obtained from the NIA at 19 months of age and housed at the Harvard T.H. Chan School of Public Health (Boston, MA). Mice were group housed 3-4 per cage for the duration of the study in static isolator cages at 71°F with 45–50% humidity, on a 12:12 hour light-dark (07:00am– 07:00 pm) cycle. After arrival, mice were fed a control diet (Research Diets, New Brunswick, NJ) until the start of the study. Mice were then randomized to one of three groups: *ad libitum* diet, methionine restriction (0.1% methionine) or 40% CR. CR was started in a stepwise fashion decreasing food intake by 10% per week until they reached 40% CR at week 4. CR intake was based upon *ad libitum* intake. Mice were monitored weekly for bodyweights and food intakes. Fasting blood samples (4-6h) were taken at sacrifice (after 6 months on the diet) by cardiac puncture. Approximately 200 μl of whole blood in 1 μl of 0.5 mM EDTA was collected. The tube was spun, and the plasma removed. The remaining blood pellet was frozen at −80ºC until further analysis. Custom mouse diets were formulated at Research Diets (New Brunswick, NJ; catalog #’s A17101101 and A19022001).

For our liver calorie restriction studies, male C57BL/6J wild-type mice bred in-house at Harvard Medical School were singly housed with ad libitum (AL) access to water at the New Research Building. Male C57BL/6J mice singly housed in the animal facility of the New Research building at Harvard Medical School. These mice were first fed ad libitum (AL) access to water before being switched to house chow (LabDiet® 5053), either AL or 30% CR starting at 3-months old. Food intake was gradually reduced 10% per week for CR mice and body weight as well as food intake were monitored weekly. For both treatment groups of mice, food was placed on the floor of the cage each day at 8:00 a.m. ± 1 hour. Livers were collected after approximately 10-months of treatment.

For our studies of high-fat diet in liver, mice were housed 3-4 animals per cage with access to water in the New Research Building animal facility at Harvard Medical School. Approximately 3-4 months before the experiment, mice were switched to house chow (LabDiet® 5053). Next, a subset of mice were switched to high-fat AIN-93G diet (HFD) (modified by adding hydrogenated coconut oil to provide 60% of calories from fat, HC) starting at approximately 5 months of age for the remainder of their lives. Livers were collected from mice after 10-months of treatment.

To assess the effect of partial reprogramming in mouse liver, C57BL/6 wild-type mice were ordered from Charles River Laboratories. After being acclimated in the housing facility for at least 1 week, the mice were injected with AAV.PHP.eB viruses via the retro-orbital route to express either GFP or OSK in the liver^57^. A Tet-Off system was used to control the expression of GFP and OSK. Specifically, AAV encoding TRE-OSK was co-injected with AAV encoding CMV-tTA, and AAV encoding TRE-GFP was co-injected with AAV encoding CMV-tTA to express either OSK or GFP. One month after the AAV injection, the mice were euthanized, and the liver tissues were collected for genomic DNA extraction for the TIME-Seq experiment.

Liver DNA from intervention samples was prepared as detailed above using the Chemagic 360 system for DNA extraction and our partially automated DNA normalization protocol. Normalized DNA was prepared with TIME-Seq using mouse clock enrichment baits.

### Cell culture time-course and DNA extraction

Five independent cell lines of low-passage mouse embryonic fibroblasts (MEFs) derived from C57BL/6 mice were thawed and cultured in low oxygen conditions (3% v/v) in DMEM with 17% FBS (Seradigm) and 1% penicillin/streptomycin plus 3.8 μL of β-mercaptoethanol per bottle of DMEM (500 mL). Likewise, five cell lines of low-passage adult ear fibroblasts were cultured with DMEM plus 10% FBS, 1% penicillin/streptomycin, and 3.6 μL of β-mercaptoethanol per bottle of DMEM in the same low-oxygen conditions. After thawed cell lines became confluent in a 150 mm cell culture dish, cells were split into a 12-well cell culture dish and an aliquot of ≥ 1 million cells were taken for the initial timepoint by first washing cells with cold PBS and then freezing at −80ºC. Two more aliquots of each cell line were collected over the following 28 days in the same fashion, expanding each culture to 100 mm dish and collecting cell lines simultaneously. DNA was extracted from cell pellets on the Chemagic 360 system (Perkin Elmer) with the 250 μL blood kit (Perkin Elmer, CMG-746) and samples were prepared with TIME-Seq.

### Human whole-blood DNA samples

The database of the Mass General Brigham Biobank, a biorepository of consented patient samples at Mass General Brigham (parent organization of Massachusetts General Hospital and Brigham and Women’s Hospital), was queried for subjects from the “Healthy Populations” cohorts for whom DNA samples were available. From this pool of subjects, samples were selected for clock development that were demographically representative of the U.S. population from ages 18-103 years old. DNA samples were obtained from the biobank and deidentified before distribution for TIME-Seq. Sample handling, data analysis, and study design were approved by the Mass General Brigham Institutional Review Board (protocol number 2021P003059). Deidentified DNA samples were split into an initial cohort for model training and testing and a validation cohort. Cohorts were separately normalized to a starting concentration of 10 ng/μL, prepared with TIME-Seq, and sequenced on an Illumina NovaSeq. Validation libraries were also shallow sequenced on a MiSeq.

## Data availability

Raw FASTQ sequencing data will be deposited to NCBI Sequence Read Archive (SRA) and organized in a BioProject.

## Code availability

Code for demultiplexing and read processing as well as analysis of clocks will be provided on GitHub at https://github.com/patricktgriffin/TIME-Seq. *scAge* prediction code is provided on GitHub at https://github.com/alex-trapp/scAge.

## Acknowledgements

We would like to thank Dr. Ryan Rogers for help editing this manuscript. We would like to thank Dr. Alex Nguyen Ba and Dr. Sarah Boswell for providing the Tn5 purification and reaction protocols, as well as Dr. Chiara Ricci-Tam for providing hybridization enrichment protocols. We would like to thank the Bauer Core Facility at Harvard for sequencing and DNA extraction help as well as the Harvard Microbiology Department for allowing us access to their Illumina MiSeq. We would also like to thank the Seidman Lab at Harvard Medical School for allowing us to use their Agilent Tapestation. This work was supported by the National Institutes of Health grant to The Jackson Laboratory Nathan Shock Center of Excellence in the Basic Biology of Aging (AG038070). P.T.G was supported by the National Science Foundation Graduate Research Fellowship (DGE1745303) and the NIH-NIA F99/K00 award (AG073499), and A.E.K was supported by a Diamond/AFAR Postdoctoral Transition Award in Aging (DIAMOND19036) and an NIA K99/R00 Fellowship (K99AG070102). N.N.H. and M.K.E. are supported by the Intramural Research Program (IRP) of the National Institute on Aging (NIA), National Institutes of Health. This work was also funded by NIH/NIA (R01AG019719 to D.A.S; P01AG055369 to S.J.M.; AG065403 and AG047200 to V.N.G) and the Glenn Foundation for Medical Research (to D.A.S.).

## Author Information

P.T.G conceived the project, designed TIME-Seq library preparations, and carried out most preliminary experiments and optimization of the TIME-Seq protocol. A.E.K organized mouse cohorts and performed frailty index and blood parameter measurement. P.T.G., A.E.K., J.L., M.A., N.H., M.S.M., A.L.M., C.C., and D.L.V. collected mouse samples extracted DNA, and normalized DNA. P.T.G., A.E.K., and J.L. performed library preparations. P.T.G. developed the bioinformatic pipeline for read processing and performed most of the bioinformatic analyses. C.K. provided the coverage-filtered RRBS methylation matrix used for TS-RRBS rDNA clock training. A.T. developed an application of *scAge* to shallow-sequenced-TIME-Seq data from both mice and humans, with input from C.K., and performed s*cAge* analyses. J.R.P. and V.N.G. obtained and de-identified human blood DNA samples, administered related data, and provided input on human clock analysis. A.T., M.V.M., and V.N.G. provided JAX mice blood samples used for clock training. N.N.H. and M.K.E. provided human DNA samples and consulted on human analysis. M.R.M., J.R.M., and S.J.M. provided dietary intervention samples.

J.A. performed treatments and provided liver samples from calorie restriction and high-fat diet mouse experiments. X.T. performed treatments and provided mouse livers from AAV-OSK and AAV-GFP control mice. P.G. completed the cell culture time course with help from J.L. D.A.S. provided feedback on project goals and directions. V.N.G. supervised the development and application of *scAge*. P.T.G. wrote the manuscript with input from D.A.S. and A.E.K. All authors edited and approved the manuscript.

## Ethics Declaration

P.T.G. and D.A.S. are named inventors on a patent application related to TIME-Seq methods filed by Harvard Medical School and licensed to Tally Health. D.A.S is a founder, investor, and equity owner for Tally Health and P.T.G. has received minor equity compensation as a consultant to Tally Health. Additional info on D.A.S. affiliations not directly related to this work can be found at sinclair.hms.harvard.edu/david-sinclairs-affiliations. A.T., C.K., and V.N.G. are named inventors on a patent application related to *scAge* filed by Brigham and Women’s Hospital. All other authors have nothing to disclose.

---

**Extended Data Figure 1.**
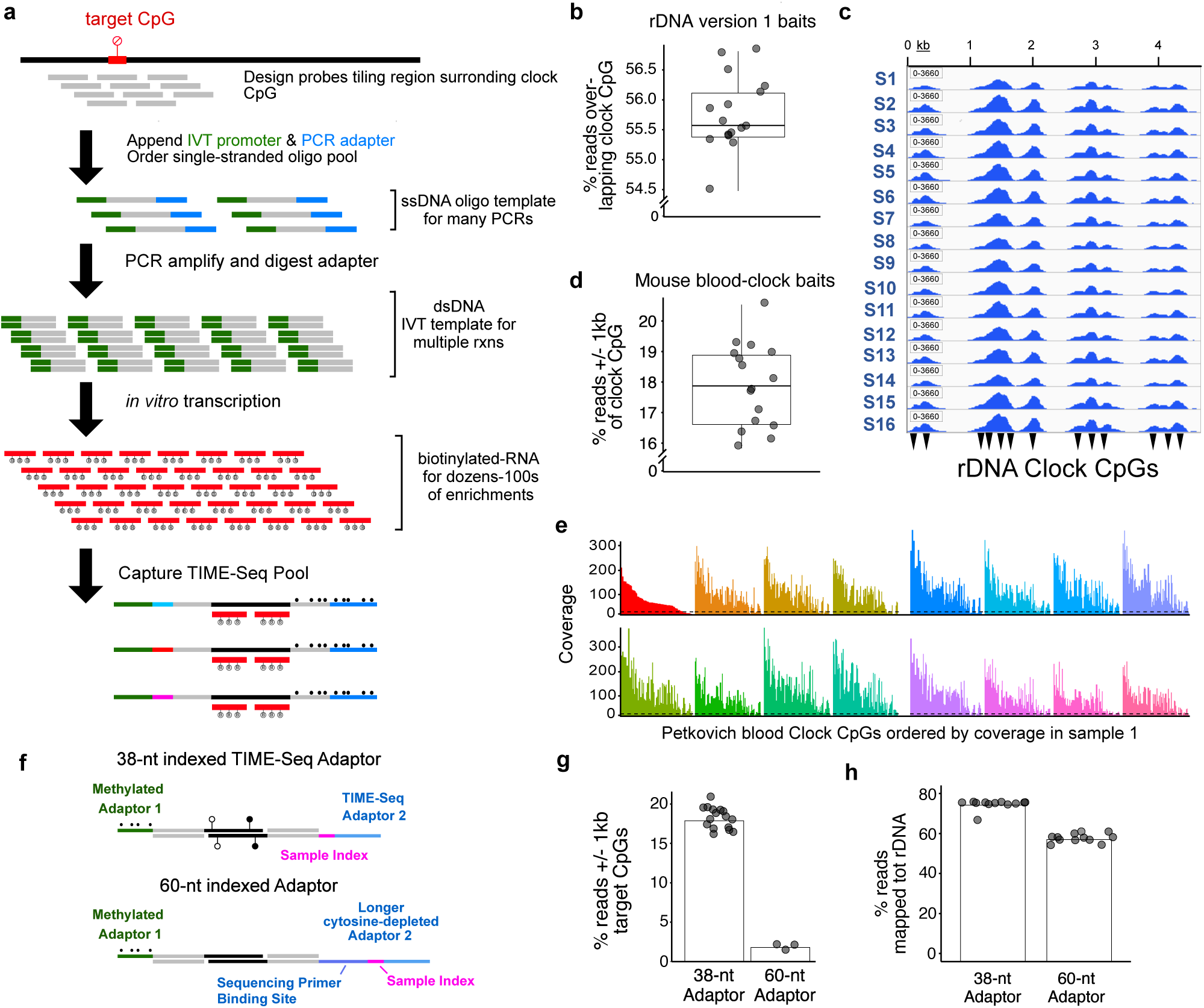
Biotinylated-RNA bait production and initial hybridization enrichment testing. **a**, Schematic of steps involved in production of biotin-RNA baits from single-stranded oligo pools for target enrichment in TIME-Seq libraries. The percent of reads overlapping target RRBS mouse rDNA clock CpGs (**b**) and an IGV browser screenshot of mapped-read pileups (**c**) using version 1 rDNA baits for enrichment of a TIME-Seq pool. Reads on-target (**d**) and mouse RRBS blood clock (Petkovich et al., 2017) CpG coverage (**e**) using mouse-blood specific baits in a pilot experiment targeting non-repetitive clock loci. Dotted line represents coverage cut-off of 10. Pools in both rDNA and blood clock pilot enrichments were sequenced with approximately 1 million paired end (PE) reads each in pools of 16 samples. **f**, Adaptor design schematic for comparison of TIME-Seq adaptors with longer barcoded adaptors. Comparison of on-target reads in short TIME-Seq and long cytosine-depleted adaptor designs for both mouse blood clock (**g**) and (**h**) rDNA (version 1) baits enrichments.

**Extended Data Figure 2.**
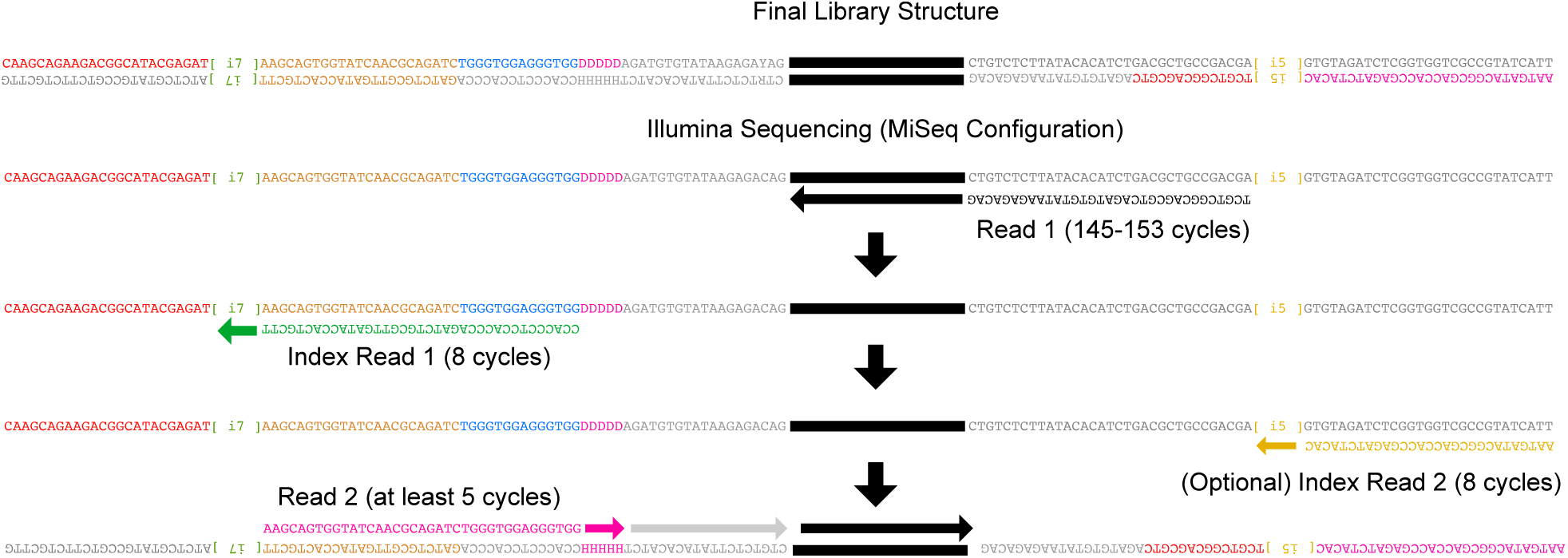
TIME-Seq library and sequencing schematic. Schematic representation of final library structure (top) and Illumina sequencing (bottom) steps required to sequence TIME-Seq libraries. Index read 1 and Read 2 primers are custom primers.

**Extended Data Figure 3.**
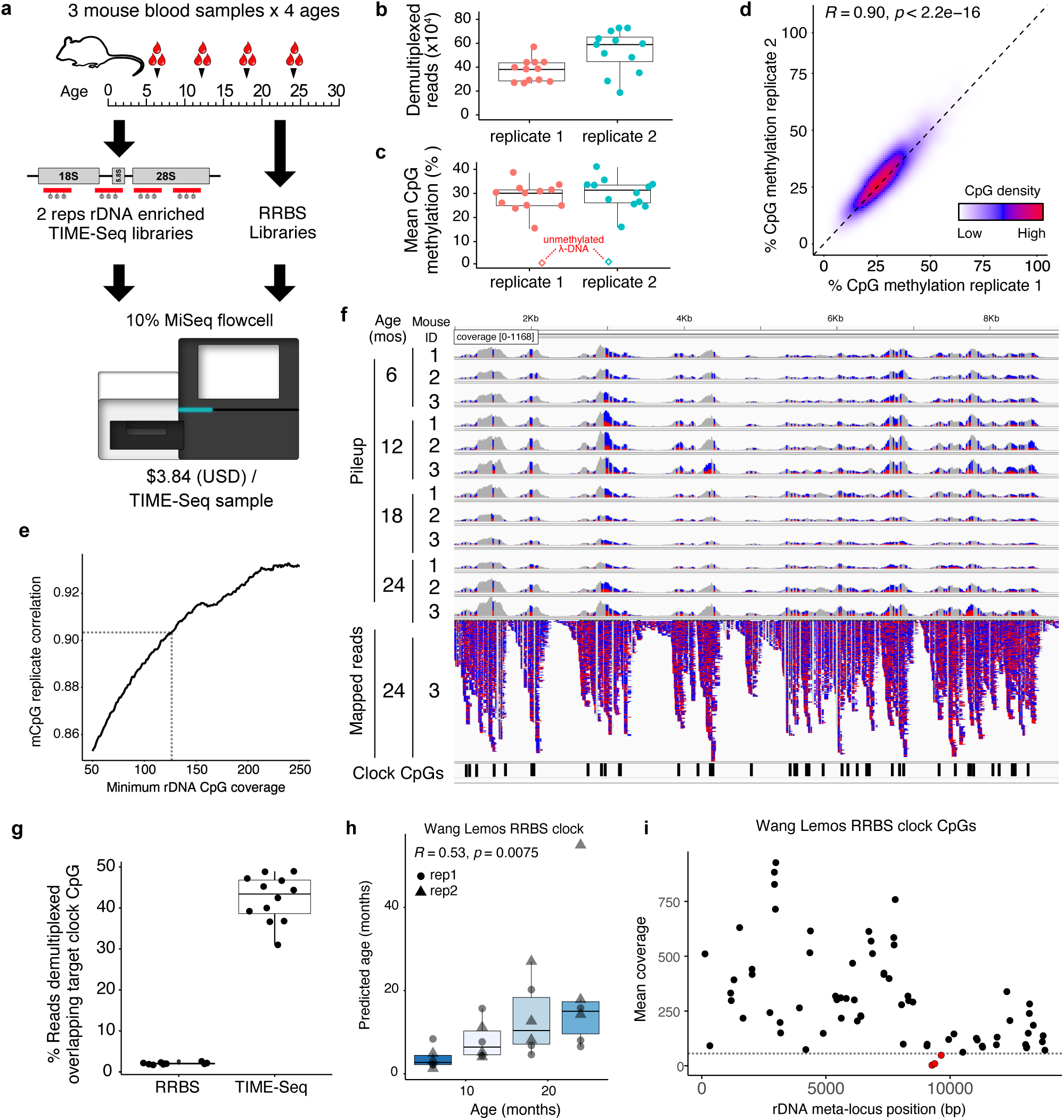
Small pilot-experiment sample metrics, correlation of rDNA CpG methylation, and age predictions using a reported RRBS-based rDNA clock. **a**, TIME-Seq pilot experimental design using mouse blood DNA from 4 age groups and preparing 2 replicates of each sample with rDNA baits (version 1) as well as RRBS libraries to be sequenced as a fraction of a Illumina MiSeq sequencing run. **b**, Demultiplexed reads from TIME-Seq pools. **c**, Mean CpG methylation from reads mapped to the mouse ribosomal DNA meta-locus. Unmethylated lambda phage DNA control is represented as a diamond. **d**, Percent methylation from reads mapped to ribosomal DNA meta-locus in replicate 1 and replicate 2 in CpGs with coverage at least 125. **e**, Replicate correlation from different coverage cutoffs in the rDNA. **f**, Pileup tracks for samples from a TIME-Seq pool (replicate 1) as well as mapped reads from sample 24-3. Reads are colored by mismatch: blue for T (unmethylated) and red for C (methylated). RRBS rDNA clock coordinates are illustrated on the bottom by black rectangles. **g**, Percent of reads directly overlapping clock CpGs from TIME-Seq libraries (N=12; mean from 2 replicates) and shallow-sequenced RRBS libraries (N=10). **h**, Wang and Lemos (2019) RRBS clock predictions using TIME-Seq data enriched for clock loci. **i**, Coverage of each clock locus in the original RRBS rDNA clock. CpGs shown in red have mean coverage less than 50.

**Extended Data Figure 4.**
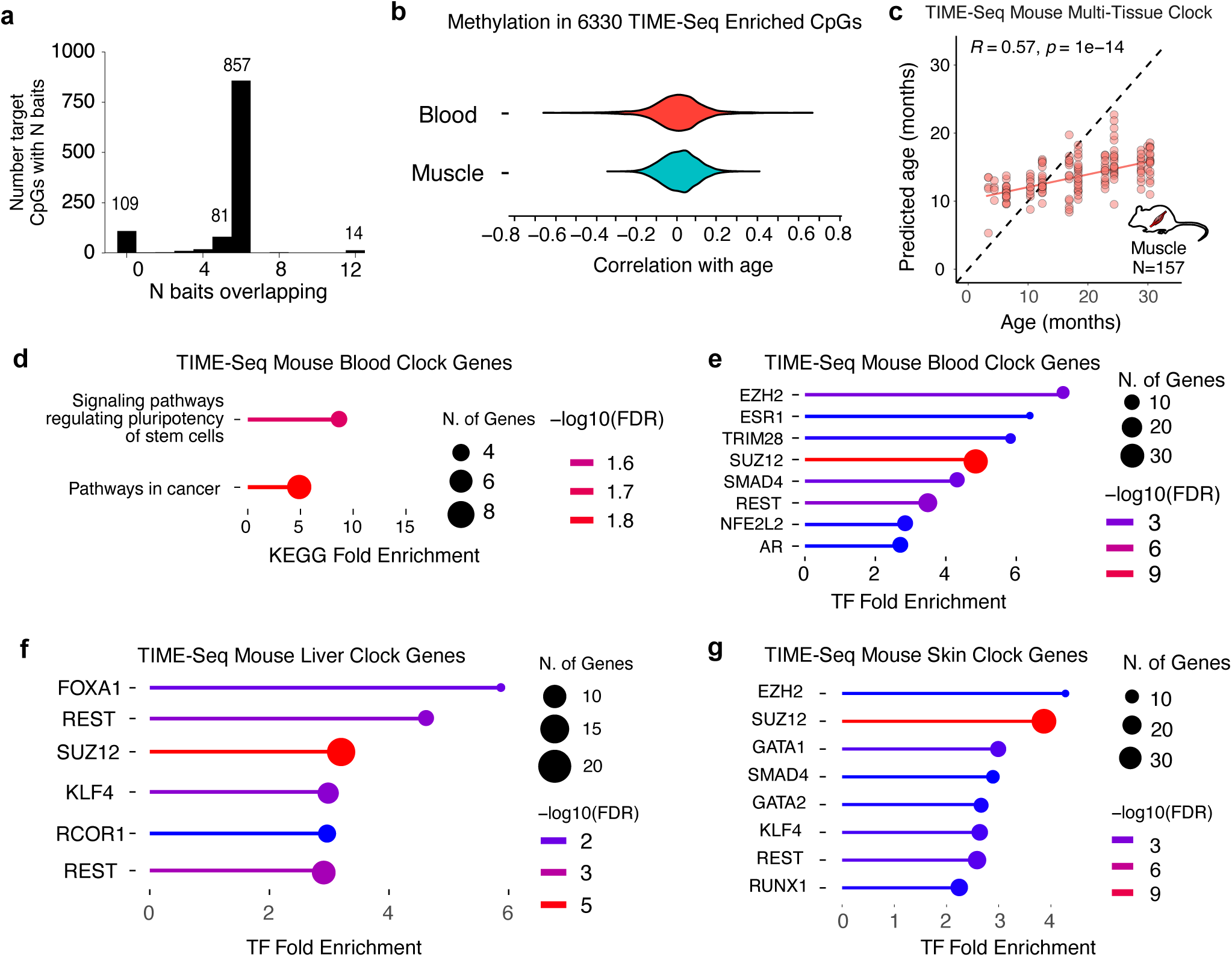
Additional data related to mouse multi-tissue and tissue-specific clock training and testing. **a**, Baits overlapping target loci used for mouse clock CpG enrichment. **b**, Comparison of correlation for blood CpGs and muscle CpGs from the enriched CpGs in TIME-Seq data. **c**, Age predictions from the TIME-Seq Mouse Multi-tissue Clock applied to the 157 mouse muscle samples. **d**, KEGG analysis of TIME-Seq Mouse Blood Clock loci. **e-g**, Transcription factor enrichment analysis for genes associated with the blood (e), liver, and skin (g) clock CpGs.

**Extended Data Figure 5.**
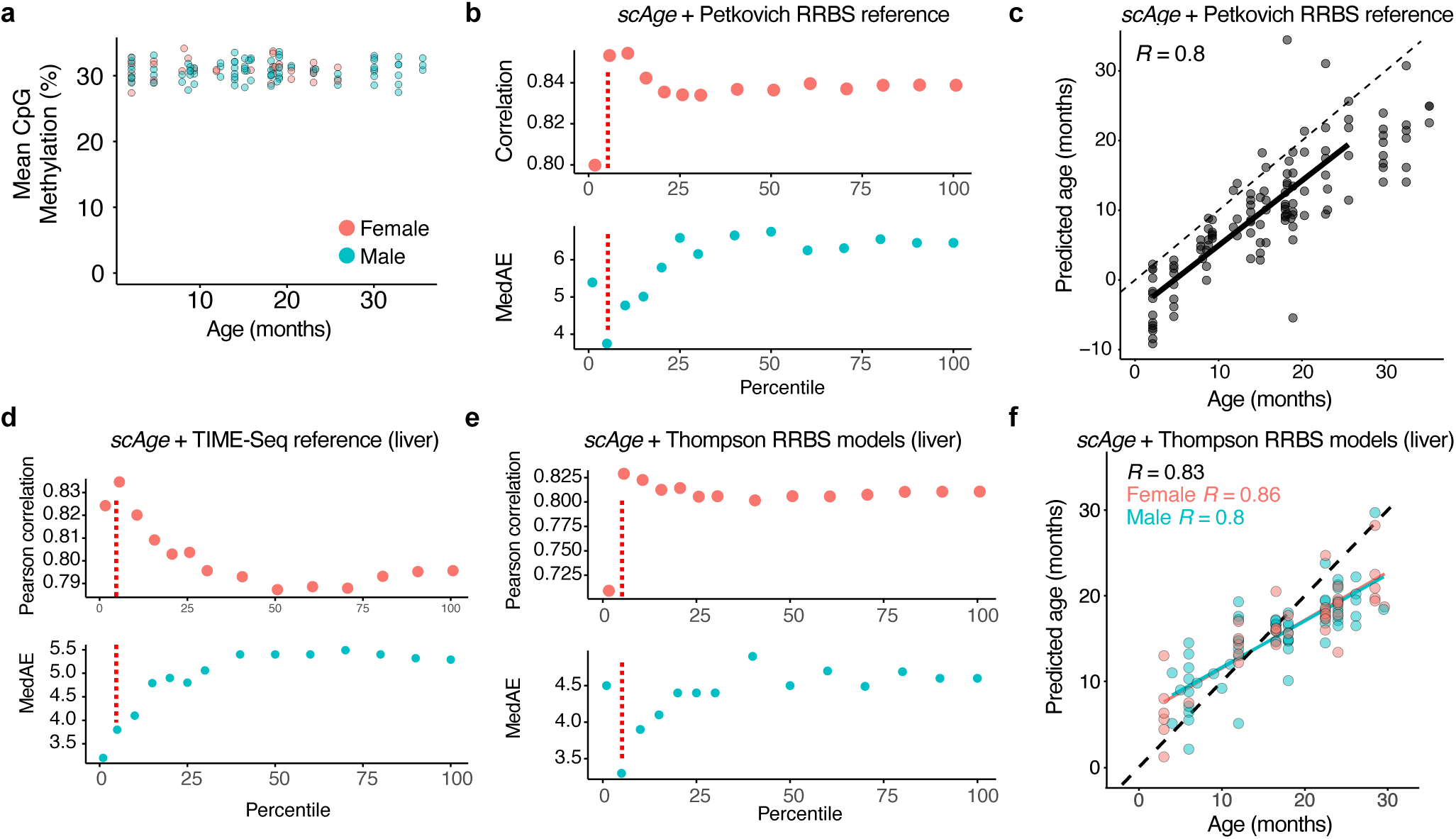
Data related to *scAge*-based shallow-sequencing predictions in TIME-Seq samples. **a**, Mean CpG methylation in shallow sequenced samples plotted against ages and colored by sex. **b**, *scAge* prediction statistics using Petkovich et al., (2017) data as model CpGs. The red line indicates the percentile chosen to be represented. **c**, Predictions in TIME-Seq samples (N=119) using the top 5% of intersecting CpGs from Petkovich et al. (2017) data as reference. Pearson correlation is shown in the top left corner. **d-e**, Prediction statistics in mouse liver samples using deep-sequenced TIME-Seq liver libraries (**d**) or RRBS liver data from Thompson et al., (2018) (**e**) as reference data. The red line indicates the percentile chosen to be represented. **f**, Age predictions in TIME-Seq liver libraries using the top 5% of CpGs from RRBS liver data as reference. Overall and sex-specific Pearson correlation coefficients are shown in the top left.

**Extended Data Figure 6.**
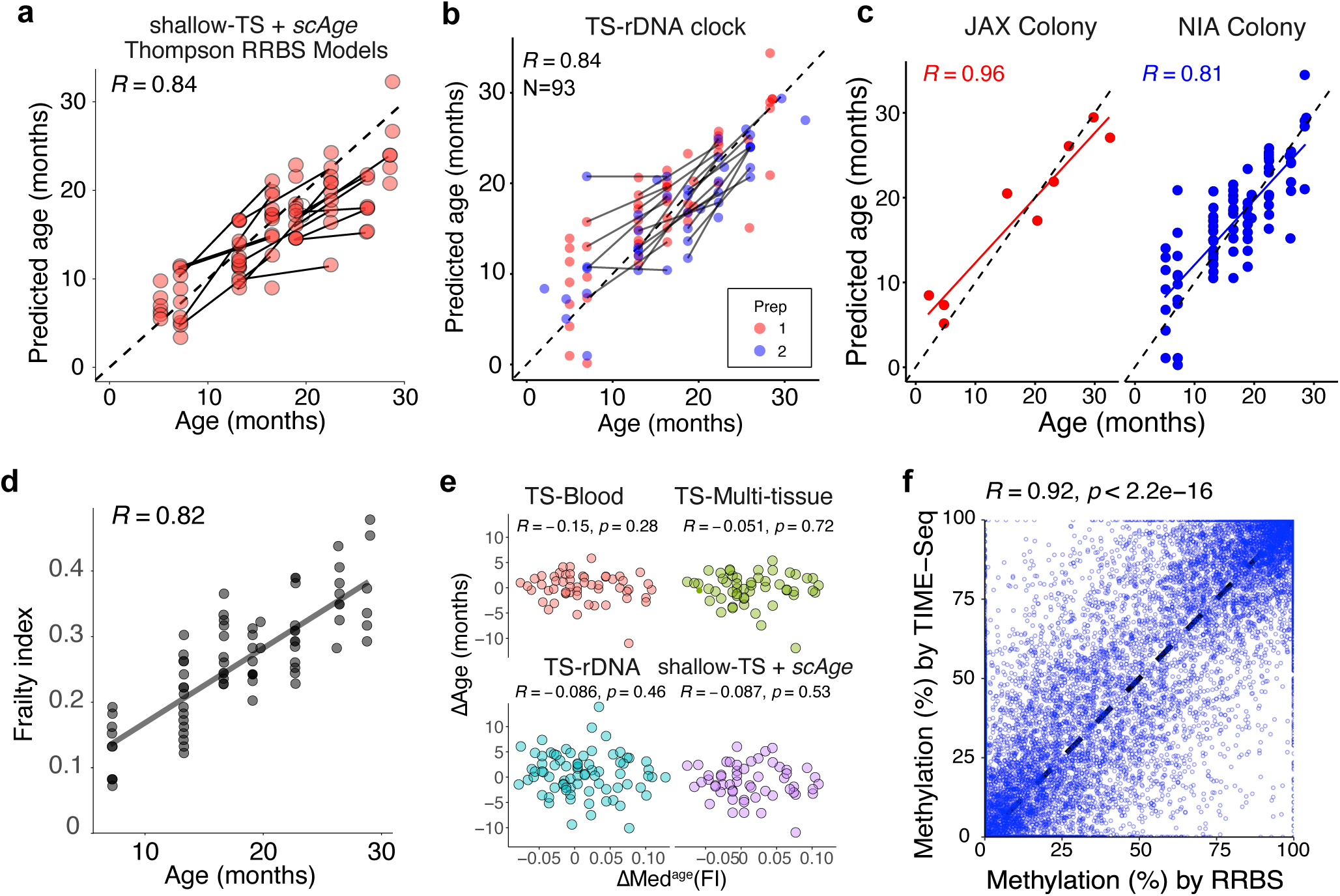
Additional Data from Validation and benchmarking of TIME-Seq. **a**, Predictions from shallow TIME-Seq data using *scAge* with the top 20% of intersecting age-associated CpGs in RRBS data as reference models. Lines connect the same mouse at different ages. **b-c**, TIME-Seq rDNA clock predictions with samples colored for (b) validation library preparation (prep) and (c) cohort of the mouse. **d**, Frailty indexes for each of the assayed mice along with Pearson correlation with age. **e**, Comparison of ΔAge and ΔMed^age^(FI) for mice in the validation cohort. **f**, Comparison between TIME-Seq CpG methylation and RRBS methylation in the same sample and CpG.

**Extended Data Figure 7.**
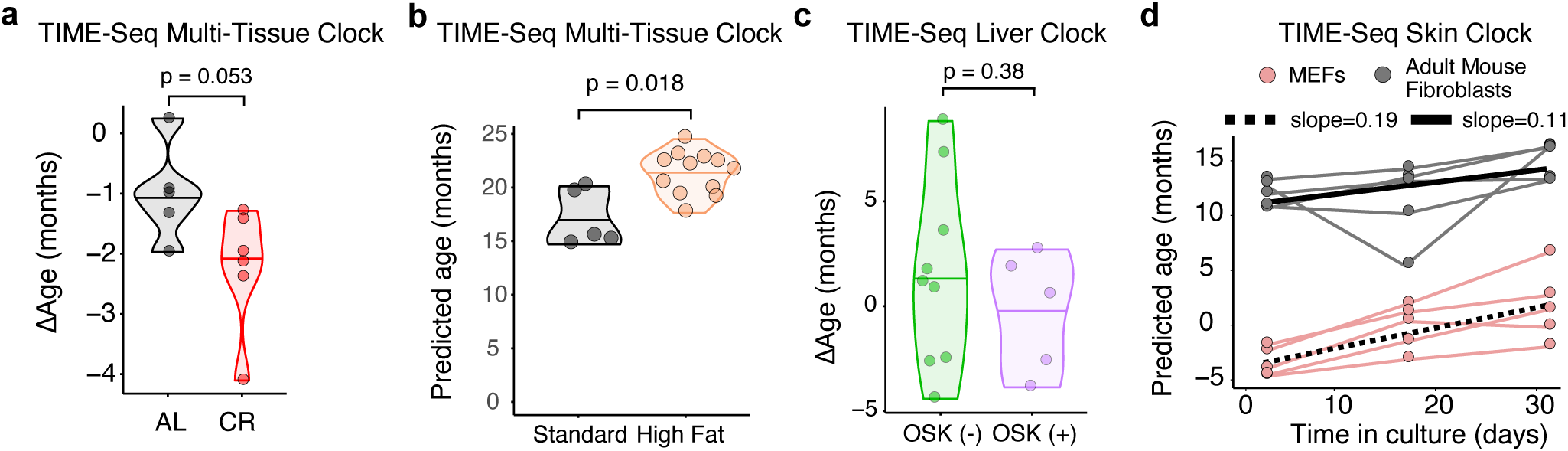
Additional Data for TIME-Seq clocks applied to intervention mice and an *in vitro* time course. **a**, Comparison TIME-Seq Multi-Tissue Clock predictions of calorie restricted mouse liver with *ad libitum* (AL). **b**, Comparison TIME-Seq Multi-Tissue Clock predictions of high fat diet mouse liver with standard diet. **c**, Comparison of TIME-Seq Liver Clock predictions in OSK-expressing, (+) OSK, and control, (-) OSK, mice. **d**, Predictions of cell culture samples using the TIME-Seq Mouse Skin Clock.

**Supplementary Table 1:**
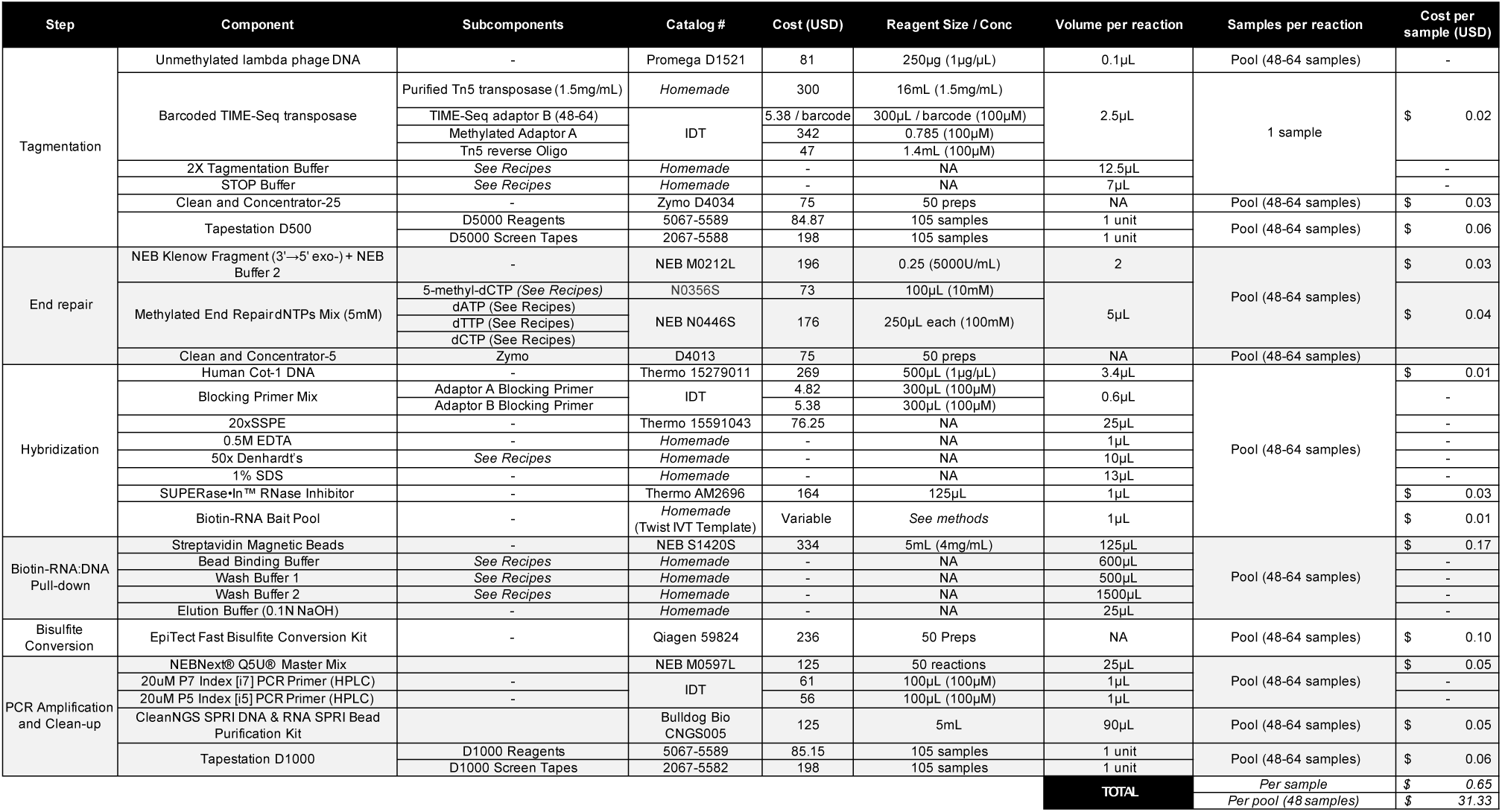
Estimated cost of reagents for TIME-Seq library preparation. Reagents that are estimated to be used in quantities that cost less than $0.01 (USD) per sample were excluded from the total cost per sample calculation.

**Supplementary Table 2:** Information on Biotinylated-RNA bait pools used for targeted enrichment of TIME-Seq libraries. * = Kb of targeted equals the target area is multiplied by copy number. Mean ≈1400 per haploid (C57BL/6 mice); mean ≈400 per haploid (Human).

| Probes | Target | # unique targets | Kb targeted | IVT Promoter |
| --- | --- | --- | --- | --- |
| Mouse ribosomal DNA v.1 baits | Wang and Lemos rDNA blood clock | 21 | 9.5* | Sp6 |
| Mouse blood-clock baits | Petkovich et al., mouse blood clock | 59 | 15.7 | T7 |
| Mouse ribosomal DNA v.2 baits | Tiling mouse rDNA meta-locus | 1 repetitive | 13.8* | Sp6 |
| Mouse discovery baits | Thompson et al., multi-tissue mouse clock.<br>Meer et al., multi-tissue mouse clock.<br>Petkovich et al., mouse blood clock | 854 | 215 | Sp6 |
| Human discovery baits | 11 previously described human clocks<br>(listed in Liu et al.) | 1289 | 324 | Sp6 |
| Human ribosomal DNA baits | Tiling human rDNA meta-locus | 1 repetitive | 13.8* | T7 |

**Supplementary Table 3:**
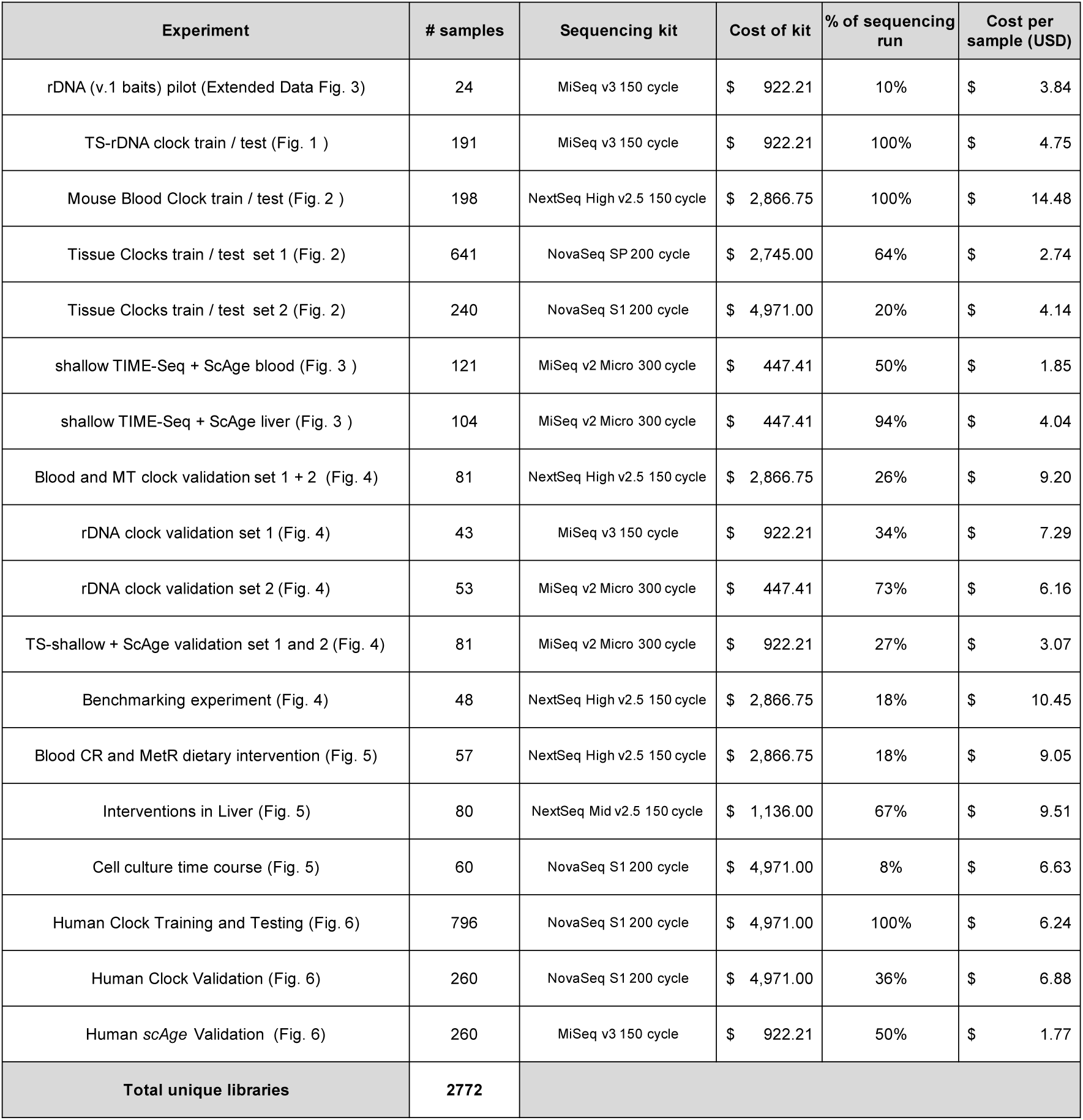
Sample number, sequencing kit, and estimated cost of sequencing for each experiment.

| Experiment | # samples | Sequencing kit | Cost of kit | % of sequencing run | Cost per sample (USD) |
| --- | --- | --- | --- | --- | --- |
| rDNA (v.1 baits) pilot (Extended Data Fig. 3) | 24 | MiSeq v3 150 cycle | \$ 922.21 | 10% | \$ 3.84 |
| TS-rDNA clock train / test (Fig. 1 ) | 191 | MiSeq v3 150 cycle | \$ 922.21 | 100% | \$ 4.75 |
| Mouse Blood Clock train / test (Fig. 2 ) | 198 | NextSeq High v2.5 150 cycle | \$ 2,866.75 | 100% | \$ 14.48 |
| Tissue Clocks train / test set 1 (Fig. 2) | 641 | NovaSeq SP 200 cycle | \$ 2,745.00 | 64% | \$ 2.74 |
| Tissue Clocks train / test set 2 (Fig. 2) | 240 | NovaSeq S1 200 cycle | \$ 4,971.00 | 20% | \$ 4.14 |
| shallow TIME-Seq + ScAge blood (Fig. 3 ) | 121 | MiSeq v2 Micro 300 cycle | \$ 447.41 | 50% | \$ 1.85 |
| shallow TIME-Seq + ScAge liver (Fig. 3 ) | 104 | MiSeq v2 Micro 300 cycle | \$ 447.41 | 94% | \$ 4.04 |
| Blood and MT clock validation set 1 + 2 (Fig. 4) | 81 | NextSeq High v2.5 150 cycle | \$ 2,866.75 | 26% | \$ 9.20 |
| rDNA clock validation set 1 (Fig. 4) | 43 | MiSeq v3 150 cycle | \$ 922.21 | 34% | \$ 7.29 |
| rDNA clock validation set 2 (Fig. 4) | 53 | MiSeq v2 Micro 300 cycle | \$ 447.41 | 73% | \$ 6.16 |
| TS-shallow + ScAge validation set 1 and 2 (Fig. 4) | 81 | MiSeq v2 Micro 300 cycle | \$ 922.21 | 27% | \$ 3.07 |
| Benchmarking experiment (Fig. 4) | 48 | NextSeq High v2.5 150 cycle | \$ 2,866.75 | 18% | \$ 10.45 |
| Blood CR and MetR dietary intervention (Fig. 5) | 57 | NextSeq High v2.5 150 cycle | \$ 2,866.75 | 18% | \$ 9.05 |
| Interventions in Liver (Fig. 5) | 80 | NextSeq Mid v2.5 150 cycle | \$ 1,136.00 | 67% | \$ 9.51 |
| Cell culture time course (Fig. 5) | 60 | NovaSeq S1 200 cycle | \$ 4,971.00 | 8% | \$ 6.63 |
| Human Clock Training and Testing (Fig. 6) | 796 | NovaSeq S1 200 cycle | \$ 4,971.00 | 100% | \$ 6.24 |
| Human Clock Validation (Fig. 6) | 260 | NovaSeq S1 200 cycle | \$ 4,971.00 | 36% | \$ 6.88 |
| Human scAge Validation (Fig. 6) | 260 | MiSeq v3 150 cycle | \$ 922.21 | 50% | \$ 1.77 |
| **Total unique libraries** | **2772** |  |  |  |  |

**Supplementary Table 4**: Estimated costs for library preparation and clock data generation. *(Attached separately)*

**Supplementary Table 5**: Oligonucleotides used for TIME-Seq library preparation. *(Attached separately)*

**Supplementary Table 6**: Clock CpG positions, coefficients, and intercepts. *(Attached separately)*

